# Early growth response 2 (EGR2) is a novel regulator of the senescence program

**DOI:** 10.1101/2020.09.30.321190

**Authors:** Eleanor J. Tyler, Ana Gutierrez del Arroyo, Ryan Wallis, Bethany Hughes, James C. Garbe, Martha R. Stampfer, Jim Koh, Robert Lowe, Michael P Philpott, Cleo L. Bishop

## Abstract

Senescence, a state of stable growth arrest, plays an important role in ageing and age-related diseases *in vivo*. Although the INK4/ARF locus is known to be essential for senescence programs, the key regulators driving *p16* and *ARF* transcription remain largely underexplored. Using siRNA screening for modulators of the p16/pRB and ARF/p53/p21 pathways in deeply senescent human mammary epithelial cells (DS HMECs) and fibroblasts (DS HMFs), we identified EGR2 as a novel regulator of senescence. EGR2 expression is up-regulated during senescence and its ablation by siRNA in DS HMECs and HMFs transiently reverses the senescent phenotype. We demonstrate that EGR2 activates the *ARF* and *p16* promoters and directly binds to the *ARF* promoter. Loss of EGR2 downregulates p16 levels and increases the pool of p16- p21- ‘reversed’ cells in the population. Moreover, EGR2 overexpression is sufficient to induce senescence. Our data suggest that EGR2 is a regulator of the p16/pRB and direct transcriptional activator of the ARF/p53/p21 pathways in senescence and a novel marker of senescence.

## Introduction

The limited replicative capacity of cultured human cells, resulting in senescence, was first described by Hayflick & Moorhead (1961) and has since been implicated to play an important role during *in vivo* ageing and age-related diseases (van Deursen, 2014). Senescence, a stable proliferative arrest, occurs in response to diverse damaging stimuli triggering up-regulation of cyclin-dependent kinase inhibitors (CDKIs), altered gene expression and subsequent nuclear and cellular morphological changes (Sharpless and Sherr, 2015). Two families of CDKIs, including p16^INK4A^ (p16) and p21^Cip1/Waf1^ (p21), can independently initiate senescence programs by directly binding and inhibiting cyclin-CDK complex phosphorylation of retinoblastoma (RB) (Dyson, 1998).

Study of *p16* regulation has revealed numerous pathways that converge to regulate *p16*, and by extension the INK4/ARF locus, which also encodes *p15*^*INK4B*^ and *p14*^*ARF*^*/p19*^*ARF*^ (*ARF*) Gil and Peters, 2006; Martin, Beach and Gil, 2014). Importantly, ARF functions to inhibit MDM2 ubiquitination and degradation of p53, leading to up-regulation of *p21*, a transcriptional target of p53. Thus, the INK4/ARF locus forms a pivotal link between the two key senescence initiation cascades (Zhang, Xiong and Yarbrough, 1998).

Epigenetic repression of the INK4/ARF locus is controlled by two polycomb repressive complexes (PRC1 and PRC2; Gil *et al*., 2004). In addition, individual transcription factors directly repress the *p16* promoter, including the hedgehog pathway component, GLI2 (Bishop *et al*., 2010), and homeobox proteins, such as HLX1, which act to recruit the PRC2 complex to the locus (Martin *et al*., 2013). Similarly, T-box proteins, TBX2 and TBX3, directly repress the *ARF* promoter (Jacobs *et al*., 2000; Brummelkamp *et al*., 2002).

Although it is well established that ETS1 mediates *p16* induction in fibroblasts by the RAS/RAF/MEK cascade during oncogenic signalling, leading to oncogene-induced senescence (Serrano *et al*., 1997), the upstream pathways activating the INK4/ARF locus in epithelial and fibroblast senescence are not well understood. To date, overexpression of the homeobox protein, MEOX2, has been identified to induce senescence in keratinocytes and fibroblasts by directly binding to and activating the *p16* promoter (Irelan *et al*., 2009), and overexpression of E2F1 induces senescence in fibroblasts via increased *ARF* expression (Dimri *et al*., 2000). However, depending on the cellular context, β-catenin can directly activate (Wassermann *et al*., 2009) or repress *p16* (Delmas et al., 2007), whilst FOXO proteins can directly activate *p15* and *ARF* (Katayama *et al*., 2008), or repress *p16* (Yalcin *et al*., 2008).

Furthermore, recent evidence has suggested that senescence is a multi-step, dynamic process throughout which the senescent phenotype evolves (Kim *et al*., 2013). Deep senescence (DS) takes over 7-10 days to develop post-senescence induction. For example, in epithelial cells, it is defined when cultures at p16-dependent stasis undergo no further expansion upon at least two serial passages (Lowe *et al*., 2015, Methods). In fibroblasts, it is further characterised by additional markers of senescence, most notably the senescence-associated secretory phenotype (SASP) (Coppé *et al*., 2008; Rodier *et al*., 2009), accompanied by elevated reactive oxygen species (ROS) levels (Passos *et al*., 2010; Lowe *et al*., 2015), and a loss of lamin B1 (Freund *et al*., 2012). Despite our growing understanding of the elaboration of the senescent state, there is a lack of knowledge of the key regulatory pathways upstream of the p16/pRB and ARF/p53/p21 pathways in DS.

We have previously demonstrated that DS is reversible in p16-positive primary adult human mammary epithelial cells (HMECs) using *p16* siRNA transfection (Lowe *et al*., 2015). Of relevance, p16-dependent epithelial senescence is independent of ARF/p53/p21 pathway activation (Garbe *et al*., 2009), whereas senescence in primary adult human fibroblasts engages both the ARF/p53/p21 and p16/pRB pathways (Alcorta *et al*., 1996; Figure 1A). We took note of previous work in human neonatal foreskin fibroblasts (HCA2) which demonstrated that p53 knockdown in senescence reinitiates DNA synthesis but with limited proliferation (Gire and Wynford-Thomas, 1998), and subsequent findings that p53 or pRB inactivation in neonatal foreskin fibroblasts (BJ), with low levels of p16, reversed senescence (Beauséjour *et al*., 2003). However, p53 inactivation or p16 shRNA knockdown followed by p53 inactivation in foetal lung WI38 fibroblasts, with higher levels of p16, did not reverse senescence, leading the authors to suggest that activation of the p16/pRB pathway may provide a dominant second barrier to senescence reversal (Beauséjour *et al*., 2003).

**Figure 1.**
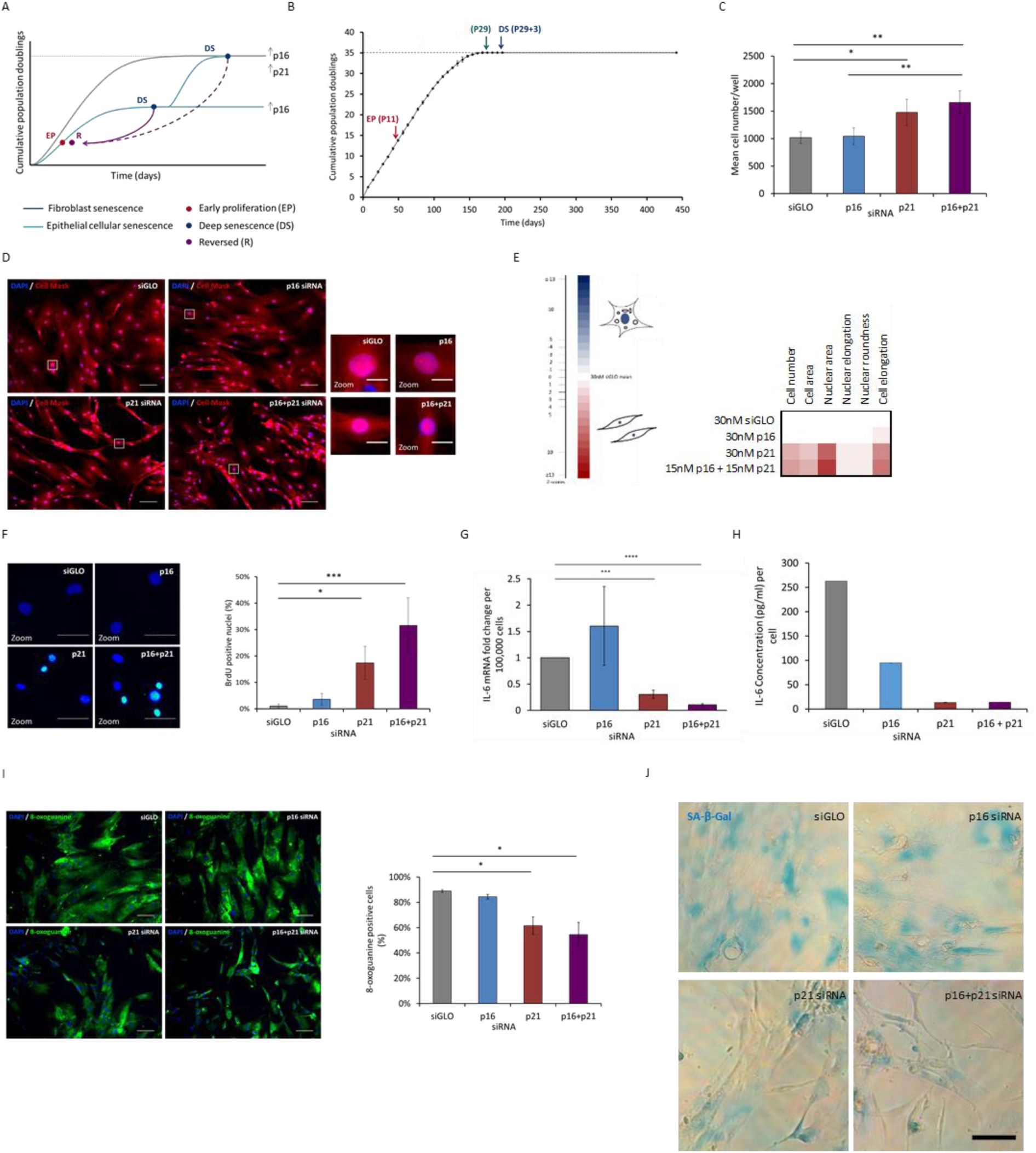
Deep senescence (DS) in primary adult human mammary fibroblasts is reversible. **(A)** Schematic illustrating epithelial and fibroblast senescence, and the DS reversal strategy. **(B)** Early proliferating (EP) fibroblasts at P11 were serially passaged until they reached senescence at P29. Deeply senescent (DS) fibroblasts were defined as a population which did not expand when kept in culture for three weeks post-senescence (P29+3). No expansion was observed in DS fibroblasts kept in culture for a further 130 days. N=1 between P4 and P6; N=2 or more between P7 and P29+3; N=1 P29+3 + 130 days. Error bars=SD of at least two independent experiments. **(C-J)** DS HMFs were forward transfected with 30nM control siRNA (siGLO), 30nM *p16* siRNA (p16), 30nM *p21* siRNA (p21), or 15nM *p16* siRNA together with 15nM *p21* siRNA (p16+p21) and fixed five days post transfection (B-E, H-I), harvested for RTqPCR at 72 hours post transfection (F) and conditioned medium collected five days post transfection (G). **(C)** DS HMFs stained with DAPI (blue) and Cell Mask (red). Size bar 100µm. Right panel=digital zoom. Size bar 20µm. **(D)** Bar chart showing mean cell number/well. * p<0.05, ** p<0.01. Error bars, SD from four independent experiments, each performed with three replicates. **(E)** Multi-parameter analysis of cellular and nuclear morphological measures. Colour coding used to illustrate the number of Z scores of the experimental siRNA value from the siGLO mean. **(F)** DS HMFs stained with DAPI (blue) and anti-BrdU (green). Size bar 50µm. Bar chart showing mean BrdU positive nuclei for each condition. * p<0.05, *** p<0.001. Error bars, SD from three independent experiments, each performed with three replicates. **(G)** RTqPCR analysis of mRNA levels of *IL-6* in DS HMFs *** p<0.001, **** p<0.0001. Error bars, SD from two independent experiments, each performed with two replicates. **(H)** Representative ELISA of secreted IL-6 levels in DS HMFs. **(I)** DS HMFs stained with DAPI (blue) and anti-8-oxoguanine (green) Size bar, 100µm. Bar chart depicting mean 8-oxoguanine positive cells for each condition. * p<0.05. Error bars, SD from two independent experiments, each performed with three replicates. **(J)** Representative images of DS HMFs stained for senescence-associated beta-galactosidase (SA-β-Gal) activity (blue). Size bar, 50µm.

Here, we show that DS in primary adult human fibroblasts with high p16 levels can be reversed using transfection of *p16* siRNA in combination with *p21* siRNA. Subsequently, we perform siRNA screens in DS HMECs and human mammary fibroblasts (HMFs) in order to further understand the key regulators upstream of the p16/pRB and ARF/p53/p21 pathways which drive senescence. In this study, we present evidence that early growth response 2 (EGR2) acts as a regulator of *p16* and transcriptional activator of *ARF* in senescence and is a novel marker of senescence.

## Results

### Reversal of deep senescence in fibroblasts

Current literature suggests that senescence is a dynamic process, and that fibroblasts in ‘light’ senescence (with low p16 levels) can be reversed, whereas DS fibroblasts (with high p16 levels), have entered a distinct, irreversible state (Beausejour, et al., 2003). As such, we began by asking whether fibroblast DS (with high p16 and p21 levels) is truly irreversible. Building on previous work in which we have reversed DS in p16-positive DS HMECs (Lowe *et al*., 2015), we hypothesised that transient knockdown using previously validated *p16* (Bishop *et al*., 2010) together with *p21* (Borgdorff *et al*., 2010) siRNAs in DS fibroblasts would induce a ‘reversed phenotype’ as characterised by a panel of senescence markers (Figure 1A).

To investigate this hypothesis, we employed senescent HMF and human dermal fibroblasts (HDFs) that had been serially passaged to senescence and cultured for a further 21 days to ensure a deeply senescent state with high p16 and p21 levels (Figure 1B, Figure S1, Methods), and developed an efficient protocol to introduce siRNA into these classically hard to transfect cells (Methods). Subsequently, we depleted *p16* and/or *p21* mRNA in DS HMFs or HDFs with potent siRNAs (Figure S2A-B), and assessed the impact on numerous cellular and molecular markers classically associated with senescence in comparison to DS cells transfected with siGLO (a negative control targeting cyclophilin B (PPIB); ‘DS+siGLO’). While depletion of *p16* with siRNA in DS HMFs (‘DS+*p16* siRNA’) did not significantly alter the arrested phenotype or cellular and molecular markers of senescence, *p21* depletion (‘DS+*p21* siRNA’) significantly increased cell number and modulated some features of senescence morphology towards an early proliferating (EP) phenotype, namely, significantly decreased cell area, nuclear area, and nuclear elongation; and significantly increased nuclear roundness and cell elongation (Figure 1B-D). Strikingly, depletion of both *p16* and *p21* in DS HMFs and HDFs (‘DS+*p16*+*p21* siRNA’) stimulated a stronger reversion to an EP morphology as characterised by multiple cellular and molecular markers (Figure 1C-E, Figure S3). Using a panel of established senescence markers, we sought to explore further the consequences of *p16* and *p21* knockdown. Quantification of proliferation using 5-bromo-2’-deoxyuridine (BrdU) incorporation confirmed the significantly increased cycling activity of ‘DS+*p16*+*p21* siRNA’ HMFs compared to ‘DS+siGLO*’* HMFs (Figure 1F). Interestingly, the percentage of BrdU positive cells in ‘DS+*p16*+*p21* siRNA’ HMFs was higher even than that observed in EP HMFs (Figure S1D), indicating that a greater proportion of the ‘DS+*p16*+*p21* siRNA’ HMFs progress through S phase during the 16-hour BrdU pulse than the EP HMFs. In agreement with the reversed phenotype, ‘DS+*p16*+*p21* siRNA’ HMFs also displayed down-regulation of the SASP proinflammatory signature in comparison to ‘DS+siGLO*’* HMFs, as illustrated by significantly decreased expression of the cytokine *IL-6* (Figure 1G) and decreased IL-6 secretion (Figure 1H). In line with the literature, IL-8 expression and secretion was also investigated but found not to be a feature of the SASP in DS HMFs (data not shown; Coppé et al., 2008). We also measured levels of 8-oxoguanine, a marker of reactive oxygen species and oxidative damage, and found a significant decrease in the ‘DS+*p16*+*p21* siRNA’ population compared to ‘DS+siGLO’ HMFs (Figure 1I). Furthermore, investigation of senescence-associated beta-galactosidase (SA-β-Gal) activity in DS HMFs following transfection, suggested a potential decrease in SA-β-Gal activity in ‘DS+*p16*+*p21* siRNA’ HMFs compared to ‘DS+siGLO’ HMFs (Figure 1J). Together, our data indicate that senescence appears to be transiently reversed in the ‘DS+*p16*+*p21* siRNA’ HMFs.

### siRNA screening reveals novel regulators of senescence

We next sought to identify novel genes that regulate the senescent phenotype. Initially, we interrogated our previously published gene expression datasets to identify genes whose expression was significantly up-regulated in HMEC DS relative to EP HMECs, and down-regulated following *p16* siRNA knockdown (Figure 2A; Lowe *et al*., 2015; GEO: GSE58035, q<0.05). In order to distinguish between the genes driving senescence and downstream ‘passenger’ genes, a siRNA screen of the top 190 genes was performed in DS HMECs (Supplementary Table 1). Each gene was targeted by a pool of three siRNAs (30nM Ambion).

To determine the effect on a panel of senescence markers for each of the 190 siRNAs, the siGLO transfected control provided a baseline for Z score generation. Using high-content analysis, 28 siRNAs (14.7%) were identified to strongly induce reversal in the DS HMECs as defined by an increase in cell number and the loss of a panel of senescence markers (i.e. mimicking the HMEC phenotype generated by *p16* siRNA). Accordingly, these 28 genes were classified as potential regulators of senescence (Figure 2B).

**Figure 2.**
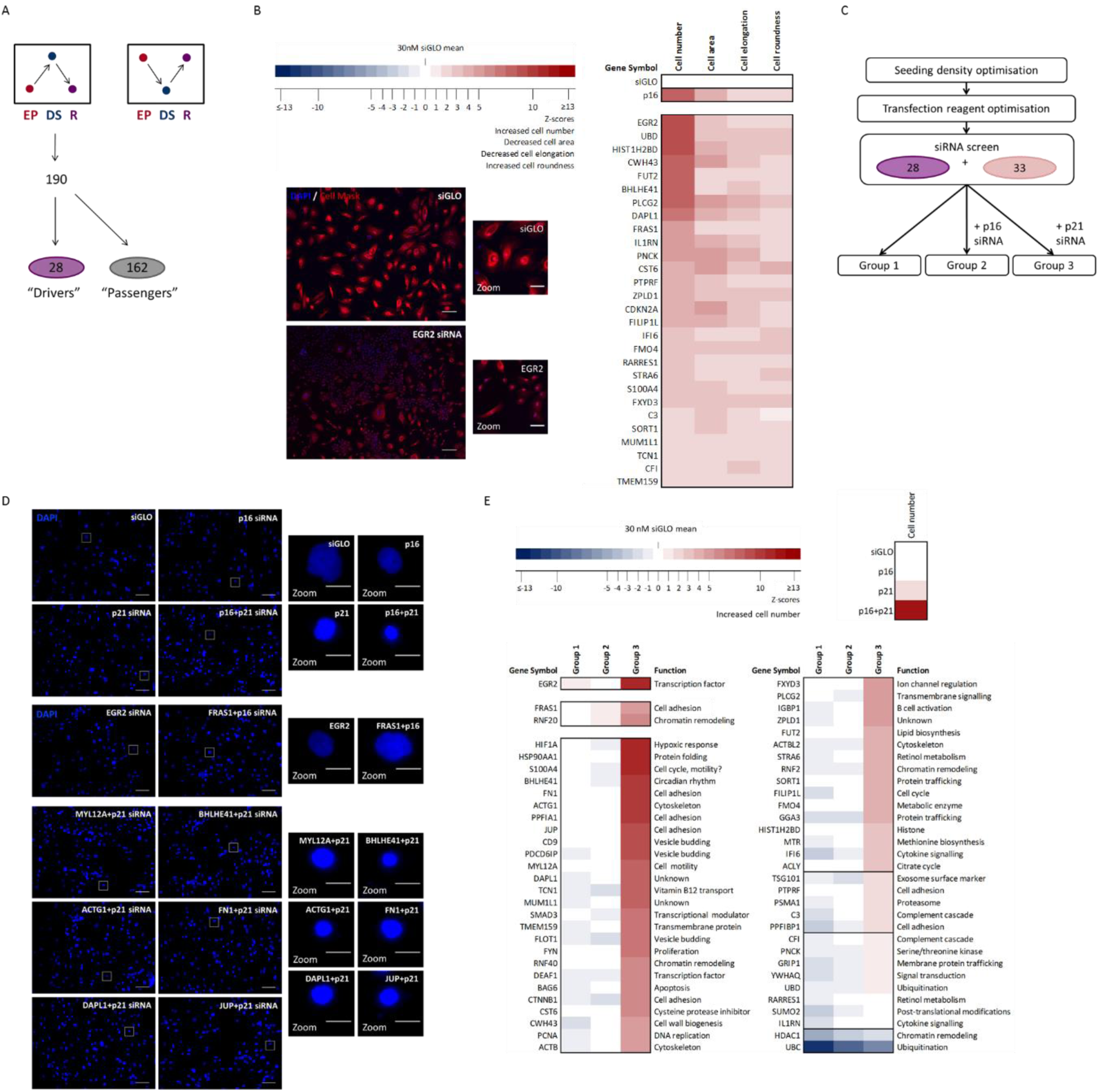
High-content screening for regulators of senescence. **(A)** Schematic illustrating mRNA microarray data which identified top 190 genes with increased expression in the deeply senescent (DS, blue) versus the early proliferating (EP, red) and reversed (R, purple) HMECs. **(B)** Results of DS HMEC screen performed twice, in triplicate. Colour coding used to illustrate the number of Z scores of the experimental siRNA value from the siGLO mean. Heatmap of Z scores for cell number, cell area, cell elongation, and cell roundness following transfection of DS HMECs. DS HMECs stained with DAPI (blue) and Cell Mask (red) following transfection with control siRNA (siGLO) or siRNAs targeting representative hit (*EGR2*). Size bar 100µm. Right panels=digital zoom. Size bar 100µm. **(C)** Schematic illustrating the experimental design of the siRNA screen. DS HMFs were forward transfected with the 60 target siRNAs in three conditions: 30nM siRNA individually (Group 1); 15nM siRNA in combination with 15nM *p16* siRNA (Group 2); and 15nM siRNA in combination with 15nM *p21* siRNA (Group 3). **(D)** DS HMFs stained with DAPI (blue) following transfection with control siRNAs (siGLO, *p16, p21, p16+p21*) or siRNAs targeting representative hits (*EGR2, MYL12A, BHLHE41, ACTG1, FN1, DAPL1, JUP*). Size bar 100µm. Right panels=digital zoom. Size bar 20µm. **(E)** DS HMF screen performed twice, in triplicate. Colour coding used to illustrate the number of Z scores of the experimental siRNA value from the siGLO mean. Heatmap of Z scores for cell number following transfection of DS HMFs with Group 1, Group 2, or Group 3 siRNAs. A brief function is assigned to each siRNA.

To further investigate the relationships between these potential 28 regulators of senescence, we constructed a protein interaction map. Briefly, these 28 genes were probed for protein interactors using the BioGRID database (Figure S4). Using Panther, KEGG pathways and Gene Ontology (GO) bioinformatics tools, 61 genes emerged (the 28 previously identified regulators, which includes p16, and 33 protein interactors) which grouped into six functional categories: immune response; cell adhesion/cytoskeleton; metabolism; transcription; growth/proliferation; and protein/vesicle trafficking (Figure S5).

We next asked whether the siRNA hits that emerged from the initial HMEC screen could also play a role in senescence in DS HMFs using this extended protein interaction network. As DS HMF reversal was found to require siRNA knockdown of both *p16* and *p21*, we hypothesised that the regulators identified in the DS HMEC screen may additionally require knockdown of either the p16/pRB or the ARF/p53/p21 pathway to induce reversal in the DS HMFs. Accordingly, DS HMFs were screened with 60 target siRNAs (27 regulators, excluding p16, and 33 interactors) in three conditions: 30nM siRNA individually (Group 1); 15nM siRNA in combination with 15nM *p16* siRNA (Group 2); or 15nM siRNA in combination with 15nM *p21* siRNA (Group 3) (Figure 2C).

Using the same approach as described for the DS HMEC siRNA screen, a hit list was generated for each of the three conditions (Groups 1, 2, and 3) (Figure 2D-E). One siRNA transfected individually (Group 1) was defined as a hit, namely early growth response 2 (*EGR2*), a transcription factor involved in several cellular processes including cell cycle and proliferation (Parkinson *et al*., 2004; Srinivasan *et al*., 2012). Two siRNAs in combination with *p16* siRNA (Group 2), fraser extracellular matrix complex subunit1 (*FRAS1*) and ring protein 20 (*RNF20*), an E3 ubiquitin ligase, were defined as hits (Figure 2D-E). Finally, 45 of the 60 siRNAs in combination with *p21* siRNA (Group 3) were defined as hits. Strikingly, eight of these 45 siRNAs induced an increase in cell number similar to the ‘DS+*p16*+*p21* siRNA’ DS HMF control, including *EGR2* and *S100A4* siRNA. As the 28 regulator siRNAs in the screen were identified as hits for senescence reversal in p16-dependent DS HMECs, it is perhaps unsurprising that 21 of these siRNAs were identified as hits requiring additional knockdown of the ARF/p53/p21 pathway to reverse senescence in DS HMFs. Furthermore, 24 of the 33 interactors investigated in this screen were also identified as Group 3 hits, highlighting the utility of the bioinformatics approach.

The top candidates from Group 1 (*EGR2*) and Group 2 (*FRAS1*), together with an additional 12 candidates from Group 3 were selected for further investigation (*HIF1A, HSP90AA1, S100A4, BHLHE41, FN1, ACTG1, PPFIA1, JUP, CD9, PDCD6IP, MYL12A*, and *DAPL1*). We performed a more detailed, independent screen with these 14 siRNAs using multi-parameter analysis of senescence-associated morphological markers with four conditions: 30nM siRNA individually (Group 1), 15nM siRNA in combination with 15nM *p16* siRNA (Group 2); or 15nM siRNA in combination with 15nM *p21* siRNA (Group 3) (Figure S6). In addition, the impact of an increased individual siRNA dose (60nM, Group 1B) was performed to identify the most potent reversed phenotype (Figure S6).

Strikingly, 11 of the 14 siRNAs transfected individually significantly decreased cell area in a dose-dependent manner (Group 1, Group 1B; Figure S7). Of these, six siRNAs transfected individually also significantly decreased nuclear area in a dose-dependent manner (Group 1, Group 1B) and *EGR2* was the only siRNA transfected individually (Group 1, Group 1B) to also significantly increase cell elongation in a dose-dependent manner. As such, EGR2 was the only siRNA that did not require knockdown of *p16* and *p21* to significantly increase cell number (Figure 2) and significantly alter three senescence-associated morphologies towards a reversed phenotype in a dose-dependent manner (Figure S7). Taken together, these data suggest that EGR2 may be acting upstream of *p16* in epithelial DS and *p16* and *p21* in fibroblast DS. To our knowledge, no direct relationship between EGR2 and senescence has previously been described, and thus we sought to explore this finding in more detail.

### EGR2 is a novel regulator of senescence

As EGR2 was identified as the top hit for reversal in both the DS HMEC and HMF screens, we next wanted to explore the role of EGR2 in senescence. First, we validated mRNA knockdown for the *EGR2* siRNA pool in DS HMFs (Figure 3A), and subsequently deconvoluted the *EGR2* siRNA pool (*EGR2 1, 2* and *3*) to determine the efficacy of each individual siRNA targeting *EGR2*. ‘*EGR2 1*’ siRNA was the least potent (Figure S8), which was subsequently reflected in the phenotype (Figure 3B-D). Using multi-parameter phenotypic analysis to control for off-target effects, we identified ‘*EGR2 3*’ siRNA as the most potent siRNA transfected individually (Figure S8) which significantly increased cell number and significantly reversed cell area, nuclear area and cell elongation (Figure 3B-D). ‘*EGR2 2*’, the second most potent siRNA transfected individually (Figure S8), produced a modest increase in cell number, and significantly reversed nuclear area and cell elongation (Figure 3B-D). Finally, the least potent siRNA, ‘*EGR2 1*’, only significantly reversed cell elongation compared to the DS+siGLO control (Figure 3D). Further characterisation of the changes to the senescence phenotype following ablation of the *EGR2* in DS HMFs revealed a significant down-regulation of the SASP factor, *IL-6*, at the transcript level (Figure 3E) and at the secreted protein level (Figure 3F). As mentioned previously, IL-8 is known not to be a feature of the DS HMF SASP (data not shown, Coppé *et al*., 2008).

**Figure 3.**
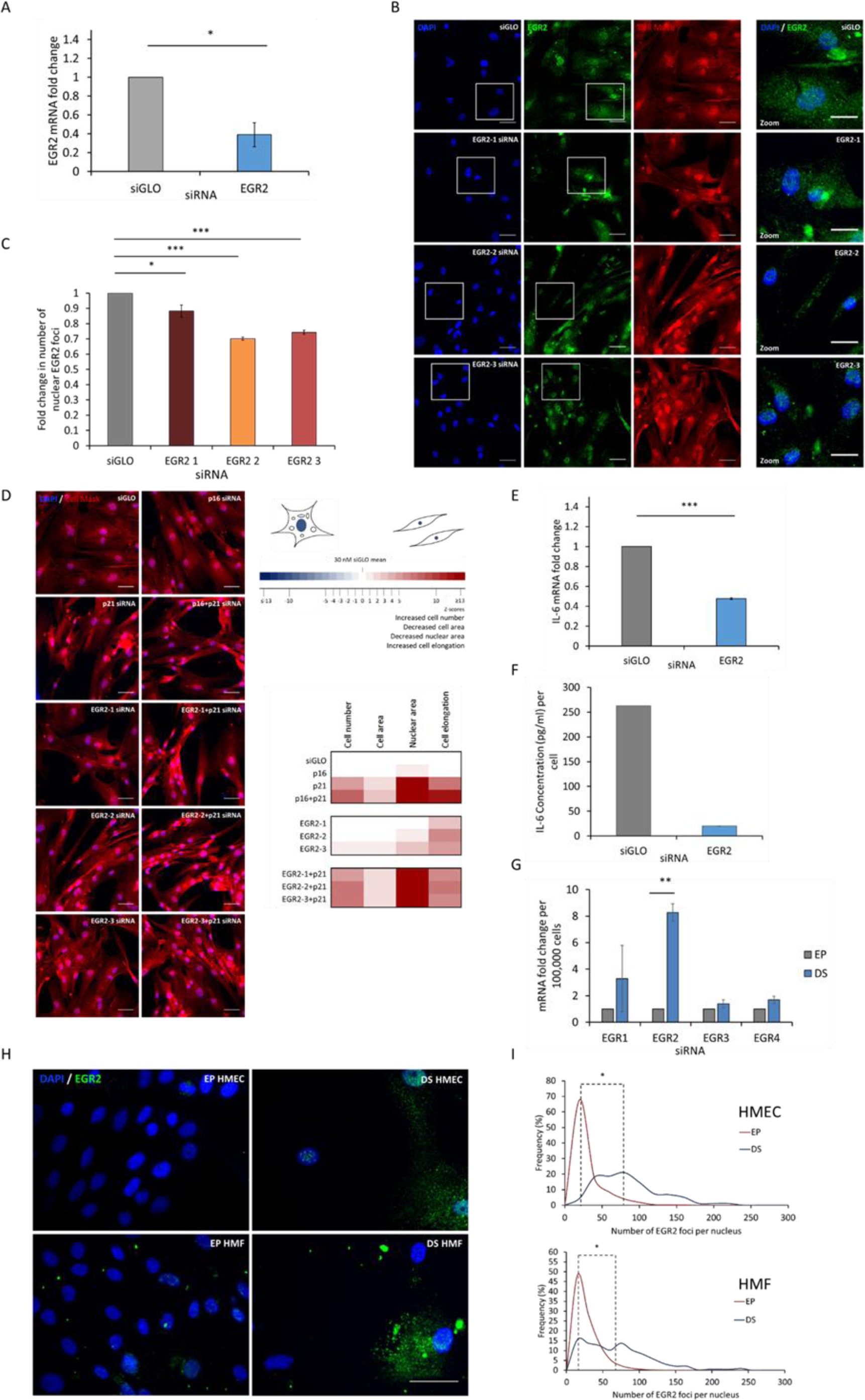
EGR2 knockdown *in vitro* reverses senescence-associated morphologies and down-regulates SASP component, IL-6, and EGR2 protein levels increase in deep epithelial and fibroblast senescence *in vitro*. **(A)** RTqPCR analysis of mRNA levels of *EGR2* in DS HMFs following siGLO or *EGR2* knockdown. ** p<0.01. Error bars, SD from two independent experiments, each performed with two replicates. **(B)** Representative immunofluorescence images of DS HMFs stained with DAPI (blue), EGR2 (green), and Cell Mask (red) following transfection with siGLO or deconvoluted *EGR2* siRNAs at 5 days post-transfection. Size bar 50µm. Right panel=digital zoom. Size bar 30µm. **(C)** Bar chart depicting median EGR2 nuclear foci following siGLO or *EGR2* siRNA knockdown. * p<0.05, ** p<0.01, *** p<0.001. Error bars, SD from two independent experiments, each performed with three replicates. **(D)** Representative immunofluorescence images of DS HMFs stained with DAPI (blue) and Cell Mask (red) following transfection with control siRNAs (siGLO, *p16, p21, p16+p21*) or deconvoluted siRNAs targeting *EGR2*. Size bar 50µm. Heatmap depicting Z scores for phenotypic validation following *EGR2* siRNA pool deconvolution in DS HMFs. Two independent experiments were performed, each in triplicate. **(E)** RTqPCR analysis of mRNA levels of *IL-6* in DS HMFs following siGLO or *EGR2* knockdown. *** p<0.001. Error bars=SD from two independent experiments, each performed with two replicates. **(F)** Representative ELISA of secreted IL-6 levels in DS HMFs following transfection with control siRNA (30nM siGLO) or 30nM *EGR2* siRNA (EGR2). **(G)** RTqPCR analysis of mRNA levels of EGR family members (*EGR1, EGR2, EGR3, EGR4*) in EP and DS HMFs. ** p<0.01. Error bars, SD from two independent experiments, each performed with two replicates. **(H)** EP and DS HMECs and HMFs stained with DAPI (blue) and EGR2 (green). Size bar 50µm. **(I)** Frequency distributions of EGR2 nuclear foci in EP and DS HMECs and HMFs. * p<0.05. Two independent experiments, each containing three technical repeats were performed.

It is important to note that the human genome encodes four EGR transcription factors, EGR1-4, that share three highly homologous DNA binding zinc finger domains that can bind to the same GC-rich consensus DNA binding motif (Beckmann and Wilce, 1997). In addition, a role for EGR1 has previously been implicated in RAF-induced oncogene-induced senescence (OIS) of human BJ fibroblasts (Carvalho *et al*., 2019) and replicative senescence (RS) of murine embryonic fibroblasts (Krones-Herzig, Adamson and Mercola, 2003). As such, we wanted to investigate the expression of EGR family members in HMEC epithelial senescence and HMF senescence. EGR2 was the only member of the EGR family with significantly increased gene expression in DS compared to EP HMECs, and EGR2 was the only member of the EGR family whose gene expression significantly decreased in reversed HMECs (GEO: GSE58035). Furthermore, investigation of EGR family member expression levels in EP and DS HMFs revealed a significant increase in EGR2, but not EGR1, EGR3 or EGR4 expression levels (Figure 3G). Collectively, these data suggest that EGR2 might be the key EGR family member acting to regulate senescence in HMECs and HMFs.

Subsequently, to further explore whether EGR2 activity and regulation is conserved across multiple senescence models and occurs *in vivo* in human tissues, we performed datamining of existing GEO datasets for HDF RS, bleomycin-induced stress-induced premature senescence (SIPS) in BJ foreskin fibroblasts, and RAS oncogene-induced senescence (OIS) in WI38 lung fibroblasts (Martínez-Zamudio *et al*., 2020) *in vitro*, as well as human skin and whole-blood with age *in vivo* (STAR Methods). The abundance of EGR2 increased during senescence across all three senescence models (Figure S9A, p<0.05). Importantly, EGR2 expression increased *in vivo* in aged human skin. In addition, a recent whole-blood gene expression meta-analysis looking at over 7,000 human samples showed that EGR2 expression significantly increases with age (Figure S9, p<0.01, Peters et al., 2015). Thus, increased EGR2 expression appears to be a feature of both *in vitro* senescence and *in vivo* ageing signatures.

EGR2 possesses a nuclear localisation signal and functions to regulate gene transcription within the nucleus, thus we hypothesised that functional EGR2 would be localised within the nucleus during senescence. Immunofluorescence staining in EP and DS HMECs revealed a significant increase of nuclear EGR2 foci in DS HMECs compared to the EP population, and in DS HMFs compared to EP HMFs (Figure 3H-I). Further investigation of EGR2 levels in a third model of senescence, oncogene-induced senescence (OIS) in IMR90 lung fibroblasts (Figure S9B), identified a significant increase in nuclear EGR2 foci in OIS fibroblasts compared to the vector control (Figure S9C). These findings support our previous mining of mRNA datasets and show that an increase in EGR2 is also observed at the protein level with the expected subcellular localisation (Figure 3H-I), thus identifying EGR2 as a novel marker of senescence in both DS HMECs, HMFs, and OIS IMR90 fibroblasts.

Finally, to explore the potential mechanisms through which EGR2 may be driving senescence and identify a panel of genes that might be regulated by EGR2 during senescence, we asked if genes identified to be up-regulated in senescence in the HMEC gene expression array were enriched for the previously published EGR2 consensus binding sequences (ACGCCCACGCA; Jolma et al., 2013; Mathelier et al., 2016) compared to randomly sampled background gene sets (Figure S10A-C). Interestingly, there was a small but significant enrichment for EGR2 binding sites at the promoters of genes up-regulated in HMEC DS. Furthermore, ten of these genes were identified as hits for senescence reversal in the DS HMEC screen, including *p16*, and nine of these were also identified as hits in the HMF siRNA screen, including the top hit *S100A4*, suggesting that EGR2 may act as a senescence regulator by activating the expression of these genes.

### EGR2 regulates senescence via the p16/pRB and ARF/p53/p21 pathways

Although previous work has identified EGR2 binding to the *p21* promoter (Srinivasan *et al*., 2012; Zheng *et al*., 2013), no investigation has yet been performed on other pathways of senescence (Figure 4J). Further examination of the INK4/ARF locus revealed previously unreported hypothetical EGR2 binding sites (ACGCCCACGCA; Jolma et al., 2013; Mathelier et al., 2016) in the *p16, p15* and *ARF* promoter regions, indicating a potential for EGR2 to bind to and regulate expression of *p16, p15* and *ARF*. As *p15* was found not to be expressed in DS HMFs (Figure S10D), we explored the potential action of EGR2 on the *p16* and *ARF* promoters. To this end, we first investigated activation of the *ARF* promoter using transiently co-transfected U2OS cells with an expression vector encoding one of each of the four members of the EGR family or E2F1, a transcription factor known to directly up-regulate ARF which acts as a positive control (Dimri *et al*., 2000), together with pGL3 luciferase reporter constructs harbouring either the promoter sequence 800bp or 3.4kb upstream of the transcriptional start site of *ARF* (pGL3 ARF 800 or plGL3 ARF 3.4, respectively, Figure 4A). Cells transfected with the pGL3 ARF 800 or with the complete ARF promoter, pGL3 ARF 3.4, displayed a significant increase in luciferase activity following transfection with the EGR2 expression vector or E2F1 positive control, but not EGR1, EGR3, or EGR4 expression vectors, thus confirming EGR2 as a direct activator of the *ARF* promoter (Figure 4A, Figure S11).

**Figure 4.**
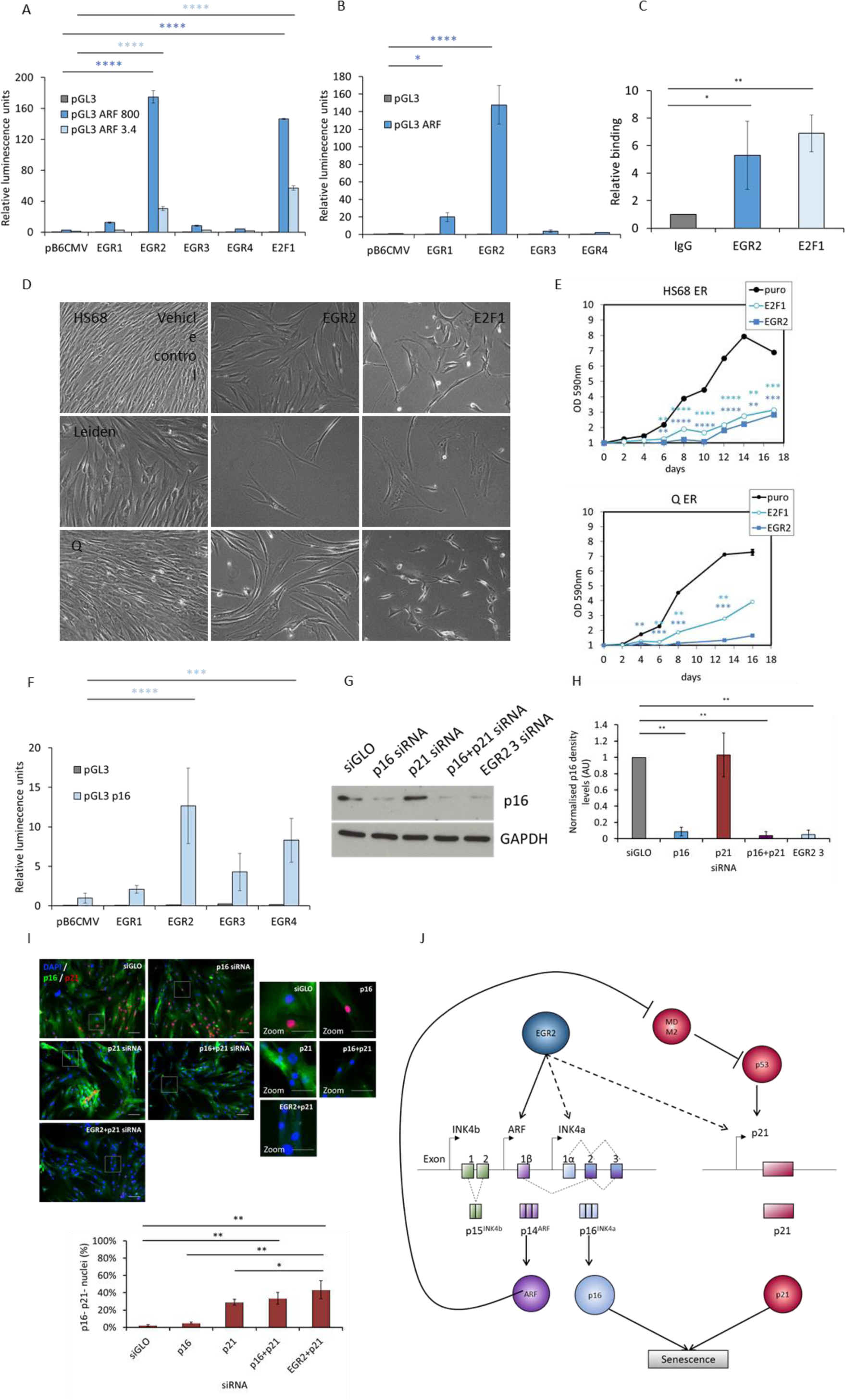
EGR2 directly binds to *ARF* and up-regulates p16 and ARF which is sufficient to induce proliferation arrest. **(A)** Mean luciferase values for activation of pGL3 luciferase reporter constructs harbouring either the promoter sequence up to 800bp or 3.4kb upstream of the transcriptional start site of *ARF* (pGL3 ARF 800 or ARF 3.4, respectively) following co-transfection of U2OS cells with expression vectors encoding each of the EGR family members (EGR1-4) or E2F1 (a positive control). Error bars, SD from two experiments. **(B)** Representative chromatin immunoprecipitation (ChIP) showing relative levels of EGR2 or E2F1 in quiescent Kit225 human T-lymphocytes or Kit225 lymphocytes following IL-2 activation. **(C)** Representative images of Hs68, p16^-/-^ Leiden or p16^+/-^ Q cells following infection with retroviral particles expressing EGR2 cDNA and selection on puromycin. Images taken at the same magnification. **(D)** Hs68 fibroblasts or p16^+/-^ Q cells were infected with retroviral particles expressing the indicated cDNAs, selected on puromycin, and assessed for proliferative capacity by periodic trypsinisation and cell counting. ** p<0.01, *** p<0.001, **** p<0.0001. Error bars=SD from three experiments. **(E)** Mean luciferase values for activation of pGL3 luciferase reporter constructs harbouring *p16* promoter sequence (pGL3 p16) following co-transfection of U2OS cells with expression vectors encoding each of the EGR family members (EGR1-4) or E2F1 (a positive control).Error bars, SD from six experiments. **(F)** Representative western blots depicting p16 levels in DS HMFs following transfection with control siRNA (siGLO), *p16* siRNA (p16), *p21* siRNA (p21), *p16* siRNA together with *p21* siRNA (p16+p21), or individual *EGR2* siRNA 3 (EGR2 3). Lysates were probed for mouse anti-p16 (JC8) and the rabbit anti-GAPDH antibody was used as a loading control. **(G)** Densitometry analysis of p16 levels in transfected DS HMFs. Analysis was performed using ImageJ software. Bars denote mean density levels. One-way ANOVA and Dunnett’s test ** p<0.01. N=2 throughout. Error bars=SD normalised to siGLO siRNA of two independent experiments. **(H)** Representative immunofluorescence images of DS HMFs stained with DAPI (blue), p16 (green), and p21 (red) following transfection with control siRNA (siGLO), *p16* siRNA (p16), *p21* siRNA (p21), *p16* siRNA together with *p21* siRNA (p16+p21), or *EGR2* siRNA together with *p21* siRNA (EGR2+p21). Size bar 100µm. Right panel=digital zoom. Size bar 50µm. Bar chart depicting mean p16 and p21 negative (p16-p21-) nuclei for transfected DS HMFs. * p<0.05, ** p<0.01. Error bars, SD from two independent experiments, each performed with three replicates. **(I)** Schematic summarising the proposed relationship between EGR2 (dark blue), ARF (purple), p16 (light blue), MDM2, p53 and p21 (red) in senescence.

Validation of the interaction between EGR2 and the *ARF* promoter was performed using chromatin immunoprecipitation (ChIP) on cross-linked DNA from quiescent interleukin 2 (IL-2) dependent Kit225 human T-lymphocytes, with low levels of *ARF* expression, and Kit225 cells following IL-2 activation which results in increased *ARF* expression (del Arroyo *et al*., 2007). Subsequently, chromatin immunoprecipitation was performed with polyclonal antibodies against EGR2, or E2F1, which acted as a positive control. Addition of IL-2 to Kit225 cells resulted in increased binding of E2F1 and EGR2 to the *ARF* promoter, demonstrating that EGR2 can be detected at the endogenous *ARF* promoter (Figure 4B).

In order to further explore the role of EGR2 in senescence, we introduced retroviral particles expressing EGR2 cDNA into normal human Hs68 diploid fibroblasts. In line with our previous observations that loss of EGR2 reverses senescence, stable overexpression of EGR2 was sufficient to induce proliferation arrest (Figure 4C-D). Interestingly, *p16*-/-Leiden cells and *p16+/-* Q cells also underwent proliferation arrest following overexpression of EGR2, indicating EGR2-mediated up-regulation of *ARF* is sufficient to induce senescence in the absence of *p16* (Figure 4C-D).

We next explored activation of the *p16* promoter and found that cells co-transfected with one of each of the four members of the EGR2 family, or E2F1, together with a pGL3 p16 construct displayed a significant increase in luciferase assay activation with the EGR2 or EGR4 expression vectors, or E2F1 positive control, confirming EGR2 and EGR4 as direct activators of the *p16* promoter (Figure 4E). As EGR4 expression is not increased in DS compared to EP HMECs or HMFs ((GEO: GSE58035, Figure 3G), we suggest that EGR2 may be important for activation of the p16 promoter in epithelial and fibroblast senescence.

If EGR2 functions to activate the *p16* promoter and up-regulate *p16* expression, we hypothesised that ablation of EGR2 in senescent cells would lead to a decrease in p16 levels. Subsequent investigation of DS HMFs transfected with an individual potent *EGR2* siRNA (‘DS+*EGR2 3* siRNA’) revealed a significant decrease in p16 protein levels compared to DS+siGLO HMFs (Figure 4F-G). Interestingly, the level of p16 in DS+*EGR2 3* siRNA HMFs was similar to DS+*p16+p21* siRNA HMFs, indicating a down-regulation of p16 in DS+*EGR2 3* siRNA HMFs comparable to reversed HMFs (Figure 4F-G). Using immunofluorescence staining and high content analysis, we further examined p16 and p21 on a cellular level and found a significant increase in the proportion of double negative (p16-p21-) ‘reversed’ cells in DS HMFs following EGR2 knockdown in combination with *p21* siRNA (‘DS+*EGR2*+*p21* siRNA) compared to the DS+*p21* siRNA HMFs, an increase similar to that seen in the reversed DS+*p16*+p21 HMFs (Figure 4H). Taken together, these data indicate that EGR2 functions to up-regulate *p16* and *ARF* expression in senescence which is sufficient to induce proliferation arrest, demonstrating that EGR2 acts as a novel regulator upstream of p16/pRB and transcriptional activator of ARF/p53/p21 pathways in senescence (Figure 4I).

## Discussion

Here, we show that DS can be transiently reversed in human fibroblasts using *p16* siRNA in combination with *p21* siRNA transfection, as characterised by the loss of a panel of senescence markers. It is important to note here that we have shown that siRNA mediated reversal of DS HMFs is transient, with population growth slowing and cells reverting to a senescence morphology by seven days post-transfection. Further investigation is required to assess the effect of long-term, stable knockdown on DS cells, including the impact on DNA damage and telomeres. However, as previous work in our group demonstrated that *p16* siRNA knockdown can reverse DS HMECs, the discovery that *p16*+*p21* siRNA knockdown can transiently reverse DS HMFs provided a unique opportunity for uncovering novel senescence regulators in epithelial and fibroblast DS. Using siRNA screening, we identified novel regulators of senescence in HMECs and HMFs, including the transcription factor *EGR2*, extracellular matrix protein *FRAS1*, E3 ubiquitin ligase *RNF20*, and calcium-binding protein *S100A4*. Further investigation of the top hit, *EGR2*, revealed that *EGR2* ablation enables resumption of the cell cycle, reversed senescence-associated morphologies and decreased expression and secretion of the SASP component, IL-6. We demonstrate that EGR2 accumulates during *in vitro* senescence in DS HMECs,DS HMFs, and OIS IMR90 lung fibroblasts. Furthermore, we re-mined existing datasets to reveal an increase in EGR2 expression in RS HDFs, SIPS BJ fibroblasts, OIS WI38 fibroblasts, and in human tissue during *in vivo* ageing. As such, we have identified EGR2 as a novel marker of senescence across multiple senescence models, including p16-dependent epithelial DS, p16- and p21-dependent fibroblast DS, fibroblast RS, OIS and SIPS. Examination of genes differentially expressed in DS HMECs identified EGR2 binding sites in *p16* and nine siRNAs found to reverse DS HMEC and HMFs, including one top reversal hit in the DS HMFs, *S100A4*. Further investigation of the INK4/ARF locus revealed previously unreported EGR2 binding sites in all the *p16, p15* and *ARF* promoters. In support of this, we demonstrated that EGR2 activates the *p16* and *ARF* promoters and that EGR2 directly binds to the *ARF* promoter. Furthermore, stable EGR2 overexpression was sufficient to induce proliferation arrest in the presence or absence of p16. Lastly, we observed a decrease in p16 protein levels in DS HMFs following EGR2 knockdown and an increase in the p16-p21-double negative subpopulation in DS HMFs following EGR2 and p21 knockdown. Given that EGR2 overexpression activates the p16 promoter, and that silencing EGR2 downregulates p16 protein levels in DS HMFs, it is likely that EGR2 directly binds to the p16 promoter. Unfortunately, discontinuation of ChIP-quality EGR2 antibodies precluded further experiments investigating this in DS cells. Further studies are needed to confirm the direct role of EGR2 activating p16 expression.

Mutations in EGR2 have been identified to lead to inherited peripheral neuropathies, including Charcot-Marie-Tooth Type 1 (Šafka Brožková *et al*., 2012), a demyelinating form associated with dysregulated Schwann cell proliferation and cell-cycle exit (Atanasoski *et al*., 2006). Accumulating evidence indicates that EGR2, a transcription factor, plays the role of regulator in these processes (Topilko *et al*., 1994; Zorick *et al*., 1996; Decker, 2006) and has been shown to directly bind to the p21 promoter in myelinating rat sciatic nerve (Srinivasan *et al*., 2012). In addition, a role for EGR2 as a tumour suppressor has been implicated in many tumour cell types (Unoki and Nakamura, 2003), and elevated expression of EGR2 is a favourable prognostic factor in breast cancer (TCGA, 5 year survival for high expressers = 84%; 5 year survival for low expressers = 73%; p=0.000073). Despite this, little attention has been paid to its role in senescence. In the present report, our findings indicate a functional role of EGR2 in regulation of *p16* and transcriptional activation of *ARF* in senescence.

Importantly, whilst our data demonstrates a role for EGR2 in regulation of senescence, transient EGR2 reversal in DS cells does not delineate between the activity of EGR2 in senescence onset or maintenance. Future studies using stable EGR2 knockdown prior to senescence entry should be performed in order to dissect the roles of EGR2 in the onset and/or maintenance of senescence.

### Concluding remarks

Our work adds to the growing list of pathways known to directly regulate senescence. This includes *p16* transcriptional repressors, such as homeobox protein HLX1 (Martin *et al*., 2013) and the N-terminal fragment of the GLI2 transcription factor (Bishop *et al*., 2010), as well as *p16* transcriptional activators such as ETS1 (Ohtani *et al*., 2001), and homeodomain protein MEOX2 (Irelan *et al*., 2009). Importantly, we have demonstrated that EGR2 functions as a regulator of p16/pRB and direct activator of the ARF/p53/p21 pathways, thus controlling both axes of the senescence program.

It is well established that expression of p16 increases with age in human tissues (Krishnamurthy *et al*., 2006), senescent cells accumulate in sites of age-related diseases (Naylor, Baker and van Deursen, 2012), and selective clearance of p16-positive senescent cells in mice has been shown to improve health- and lifespan (Baker *et al*., 2011, 2016). As such, regulation of the p16/pRB and ARF/p53/p21 pathways by EGR2 in senescence may play an important role in ageing and age-related diseases.

Furthermore, ten of these genes were identified as hits for senescence reversal in the DS HMEC screen, including *p16*, and nine of these were also identified as hits in the HMF siRNA screen, including the top hit *S100A4*, suggesting that EGR2 may act as a senescence regulator by activating the expression of these genes.

Interestingly, EGR2 as a transcription factor has the potential to regulate a network of genes in senescence, and nine hits which reversed DS HMECs and HMFs were identified to possess an EGR2 binding site, thus we hypothesise that EGR2 may potentially regulate the expression of these genes in senescence, although this has yet to be investigated further. Future exploration of the transcriptome regulated by EGR2 in senescence could provide new insights into regulation of the senescence program and potentially identify essential senescence mediators, which could be exploited to eliminate senescent cells. As implications for senescence have been described *in vivo* for organismal ageing and age-related diseases, furthering our understanding of this network in senescence could enable identification of therapeutic targets for treatment of ageing and age-related diseases.

## Experimental Procedures

### Cells and reagents

Normal finite life-span HMECs and HMFs were obtained from reduction mammoplasty tissues of a 21-year-old individual, specimen 184, and 16-year-old individual, specimen 48, respectively, and were cultured as previously described (Garbe *et al*., 2009). Independent HMEC cultures were serially passaged from passage 6 (P6; early proliferating, EP) until p16-dependent, p21-independent stasis. Deeply senescent cultures underwent no further expansion upon at least two further weeks in culture (DS HMECs; Romanov *et al*., 2001; Garbe *et al*., 2009; Lowe *et al*., 2015), and independent HMF cultures were serially passaged from P4 until the population reached senescence at P29. DS HMFs underwent no further expansion upon at least three further weeks in culture (P29+3). Cells were cultured at 37°C in the presence of 5% CO_2_ and atmospheric O_2_. All cells were routinely tested for mycoplasma and shown to be negative.

*IMR90 ER:STOP* (vector) or *ER:RAS* (OIS) IMR90 foetal lung fibroblasts were produced as described in (Hari *et al*., 2019) and were a kind gift provided by Juan Carlos Acosta. These were maintained in DMEM supplemented with 10% FBS and 2mM L-glutamine.

U2OS cells, primary human fibroblast strain Hs68, and Kit225 T-lymphocyte cell line were maintained as previously described (del Arroyo *et al*., 2007). Leiden and Q cells were maintained as previously described (Irelan *et al*., 2009).

### siRNA transfections

The fluorescently labelled siRNA targeting cyclophilin B (siGLO) was selected as this did not influence the phenotype of either EP or DS cultures (Figure S12). HMECs were transfected with 60nM siGLO siRNA (Dharmacon) or *p16* siRNA (Qiagen) in 384-well plates using Dharmafect 3 (Dharmacon). HMFs were transfected with 30nM siGLO siRNA or *p16* siRNA or *p21* siRNA (Dharmacon) in 384-well plates or 6-well plates using Dharmafect 2 (Dharmacon). DS+siGLO, DS+*p16* siRNA, DS+*p21* siRNA, or DS+*p16*+*p21* siRNA cells were harvested for RTqPCR, western blotting or immunofluorescence as detailed below.

### Immunofluorescence

Standard fixation with 3.7% paraformaldehyde, followed by 0.1% Triton X permeabilisation and blocking with 0.25% BSA was performed prior to antibody incubations. Primary antibodies used were mouseα*p16* JC8 (1:200), mouseα8-oxoguanine (1:100, MAB3560 Millipore), rabbitα*p21* (1:1,000, 12D1 Cell Signalling), rabbitαEGR2 (1:250, H220 Santa Cruz), goatαIL-6 (1:100, AB-206-NA R&D Systems), followed by donkeyαmouse AlexaFluor-488 or goatαrabbit AlexaFluor-546 (1:500, Invitrogen), DAPI and Cell Mask Deep Red (1:10,000, Invitrogen). For 5-bromo-2’-deoxyuridine (BrdU) assays, cells were cultured in 5µM for 16h prior to fixation. An additional DNA denaturation step with 4M HCl for 10 min was performed following permeabilisation, and a conjugated mouseαBrdU-AlexaFluor-488 antibody (1:100, B35130 Invitrogen) used. Images were collected at 10X using the IN Cell 1000 microscope (GE) and the Developer analysis software (GE) was used for image analysis as described previously (Bishop *et al*., 2010).

Please also see Supporting Information.

## Acknowledgements

We thank the late Gordon Peters (CRUK) for his guidance and support.

E.J.T. was supported by the Medical Research Council (MR/K501372/1). JG, and MS are supported by U.S. Department of Energy under Contract No. DE-AC02-05CH11231 and the Congressionally Directed Medical Research Programs Breast Cancer Research Program Era of Hope Scholar Award BC141351.

## Author Contributions

Conceptualization, E.J.T., A.G.dA, C.L.B.; Methodology, E.J.T., A.G.dA, J.C.G., M.R.S., C.L.B.; Investigation, E.J.T., A.G.dA., R.W., R.L., C.L.B.; Resources, J.C.G., M.R.S., J.K.; Writing – Original Draft, E.J.T., C.L.B.; Writing – Reviewing & Editing, E.J.T., A.G.dA., R.W., J.C.G., M.R.S., J.K., R.L., M.P., C.L.B.; Supervision, M.P., C.L.B.

### Declaration of Interests

The authors declare no competing interests.

### Data availability statement

Data sharing is not applicable to this article as no new data were created or analysed in this study.

## Experimental Procedures – Supporting Information

### siRNA screening and Z score generation

In the DS HMEC screen, *p16* siRNA was amongst the 28 driver siRNAs identified to reverse HMEC p16-dependent DS (Lowe *et al*., 2015). However, in DS HMFs, *p16* siRNA alone was found to not be sufficient to reverse senescence. With this in mind, *p16* siRNA was not included as a target siRNA in the DS HMF screen, bringing the total number to 60 target siRNAs (27 drivers and 33 interactors).

**Z score** = (mean value of two independent experiments for experimental siRNA – mean value of two independent experiments for siGLO)/SD for siGLO of two independent experiments. For each of the parameters analysed, significance was defined as more than one Z score away from the siGLO mean in order to increase the window for hit detection and allow as many hits to be identified as possible. Z scores are presented as a heatmap.

### Immunoblotting and densitometry analysis

Cell were lysed in RIPA buffer supplemented with 4% protease cocktail inhibitor (Roche) and protein concentration was determined using the Bio-Rad Protein Assay kit (Bio-Rad). Lysates were re-suspended in 6X Laemmli Sample Buffer (0.1M Tris pH6.8, 20% glycerol, 1% β-mercaptoethanol, 1% sodium dodecyl sulphate (SDS), 0.01% Bromophenol blue) and used for immunodetection. Primary antibodies used were mouseα*p16* JC8 (1:1,000), rabbitα*p21* (1:1,000 12D1, Cell Signalling), rabbitαlamin B1 (ab16048 Abcam, 1:1,000) and rabbitαGAPDH (1:2,000 ab9485 Abcam). Protein separation was achieved by SDS-PAGE on 10-12% polyacrylamide gels and proteins were subsequently transferred to Hybond nitrocellulose membrane (GE) using the Bio-Rad Mini-PROTEAN III system. Membranes were blocked in 5% Milk/PBS-Tween for 1hr before overnight incubation with primary antibody at 4°C with the exception of mouseα*p16*, which was used at room temperature for 2hr and rabbitαGAPDH at room temperature for 1hr. Following 3X PBS-T washes, membranes were incubated with an appropriate horseradish peroxidase (HRP)-conjugated secondary antibody for 1hr. Bands were then visualised using Enhanced-Chemiluminescence (ECL, GE). Densitometry analysis was conducted using ImageJ software. Density levels were corrected for protein loading and were expressed relative to the negative siRNA control.

### Quantitative RT-PCR (RTqPCR)

Total RNA was isolated using Qiazol (Qiagen) according to the manufacturer’s protocol. One microgram of total RNA was reverse transcribed by the Superscript III Reverse Transcriptase (Thermo Fisher Scientific) following manufacturer’s protocol. RTqPCR reactions were performed with SYBR Green Master Mix (ABI) using a 7500 Fast Real-Time PCR System (Applied Biosystems). For siRNA knockdown experiments, RNA was extracted from DS HMFs five days post-transfection. *GAPDH* levels were quantified for each cDNA sample in separate RTqPCR reactions and were used as an endogenous control. Target gene-expression levels were quantified using target specific probes. Values were normalised to the internal *GAPDH* control and expressed relative to siGLO transfected control levels (100%). All RTqPCR reactions were run in duplicate for two independent samples.

### Enzyme-linked immunosorbent assay (ELISA)

For IL-6 analysis by ELISA, equal volumes of conditioned medium were used and assay performed as per the manufacturer’s instructions (R&D Systems, Human Il-6 DuoSet ELISA DY206). Each sample was represented twice on the plate. The absorbance readings were taken at 450nm and 570nm using a CLARIOstar Plus multi-mode plate reader (BMG Labtech). Protein concentration was then estimated according to a calibration curve obtained from the absorbance values of a dilution series of the supplied standard protein control.

### Luciferase assays

Luciferase assays were performed with pGL3-ARF-736 bp, pGL3-ARF-3.4 kb, and pGL3-p16 (generated by Eiji Hara), as previously described (Matheu, Klatt and Serrano, 2005; del Arroyo *et al*., 2007). pbabepuroEGR2 was a gift from Novartis in San Diego. pB6CMV and pB6CMVEGR1/2/3/4 were a present from FJ Rauscher lab.

### Chromatin Immunoprecipitation (ChIP)

ChIP analyses were performed with antibodies against EGR2 (C-14, sc-190) and E2F1 (H-137, sc-22820) using oligonucleotide primers specific for ARF (Forward: CCCTCGTGCTGATGCTACTG, Reverse: ACCTGGTCTTCTAGGAAGCGG), as previously described (del Arroyo *et al*., 2007).

### Database searches

Protein interaction datasets were generated using the BioGRID bioinformatics database (http://www.thebiogrid.org). These interactions were then overlaid to generate a network requiring that each interactor generated a chain with at least two other drivers, revealing a total of 33 protein interactions. Using this method, only one protein interaction network was generated. We searched KEGG pathway (http://www.genome.jp/kegg/pathway.html) and PANTHER (http://www.pantherdb.org) databases to assign functional annotations to the 28 hits that strongly induced the reversal phenotype (‘drivers’) in the DS HMEC siRNA screen as well as the 14 hits in the DS HMF siRNA screen.

### Gene Expression Omnibus (GEO) dataset mining

GEO datasets (GSE41714, GSE13330, GSE18876) available at ncbi.nlm.nih.gov/gds.

### Quantification and Statistical Analysis

An un-paired, two-tailed t-test was performed to compare the means of two groups using the Microsoft Office Excel Analysis ToolPak (Microsoft, USA). A one-way analysis of variance (ANOVA) was used to analyse the differences between the means of three or more independent groups using Prism 7 (GraphPad Software Inc., USA). A two-way ANOVA was used to analyse the differences between multiple subgroups within multiple independent groups using Prism 7. Post-hoc statistical analysis was performed using either a Dunnett’s or Tukey’s multiple comparison test. The Dunnett’s test was used to compare every mean to a control mean, whereas the Tukey test was used to compare every mean with every other mean.

**Figure S1.**
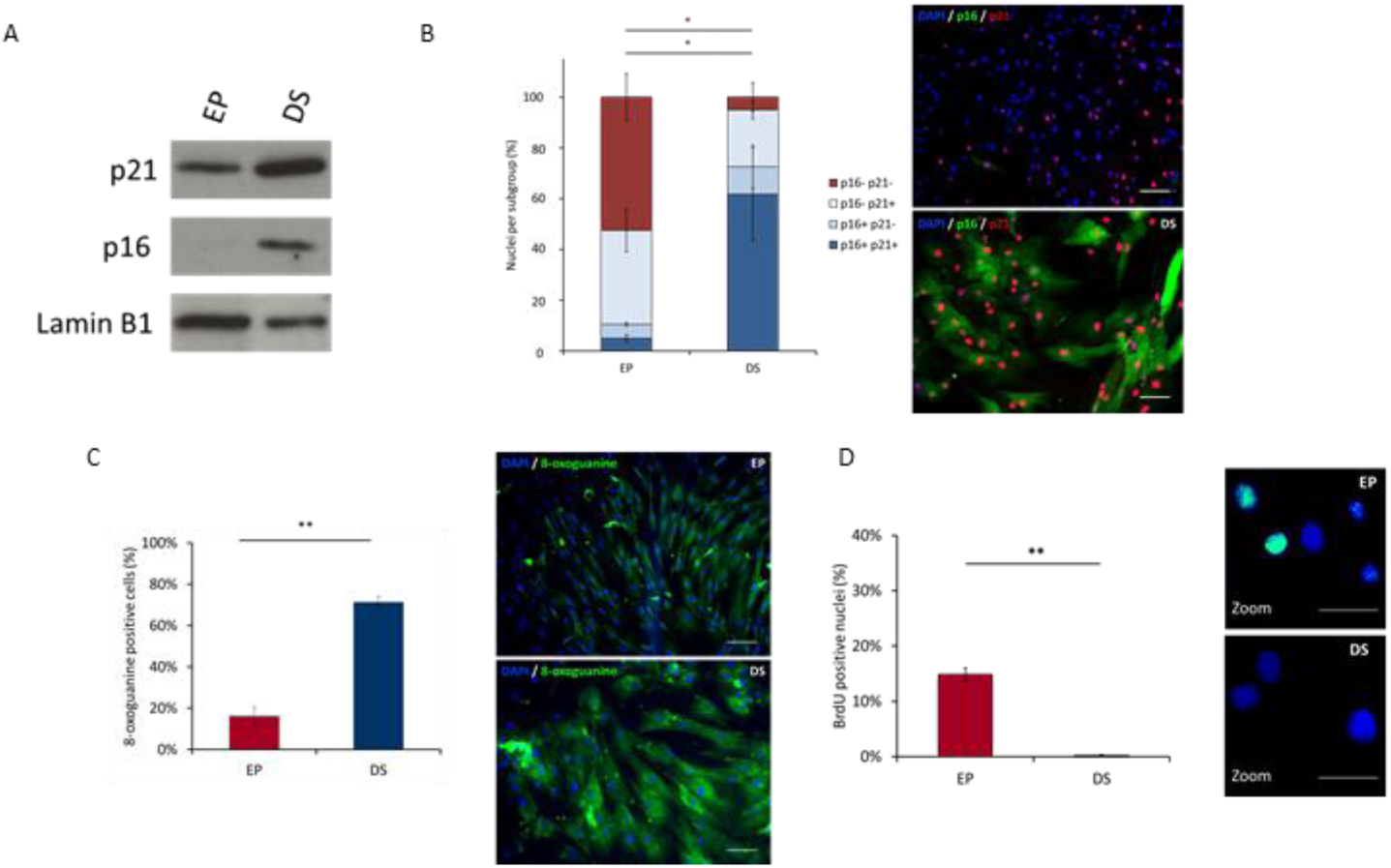
Further validation of deep senescence with an extended panel of markers. **(A-D)** EP HMFs were seeded at 10,000 cells/cm^2^ and DS HMFs were seeded at 15,000 cells/cm^2^ in 96-well plate format or 384-well plate format and harvested after five days for western blot analysis (B) or immunofluorescence (D-E). **(A)** Western blot depicting p21, p16, and lamin B1 levels in EP and DS HMFs loaded by equal cell number. Lysates were probed for rabbit anti-p21 (12D1), mouse anti-p16 (JC8), and the rabbit anti-lamin B1 antibody. **(B)** EP and DS cells were stained with DAPI, mouse anti-p16 (JC8), rabbit anti-p21 (12D1), donkey anti-mouse Alexa Fluor 488, goat anti-rabbit Alexa Fluor 546, and nuclear intensities were quantitated. Nuclear intensity thresholds were established for p16 and p21 to define positive or negative nuclei. Nuclei were classified into four subgroups: p16 and p21 negative (p16-p21-); p16 negative and p21 positive (p16-p21+); p16 positive and p21 negative (p16+ p21-); and p16 and p21 positive (p16+ p21+). Bars denote mean percentage of nuclei per subgroup. Two-way ANOVA and Tukey’s test * p<0.05. Significance colours match nuclei subgroup. N=2 throughout. Error bars, SD of two independent experiments, each performed with three replicates. Representative immunofluorescence images of EP and DS fibroblasts. DAPI (blue), p16 (green), p21 (red). Size bar, 100μm. **(C)** EP and DS cells were stained with DAPI, mouse anti-8-oxoG, donkey anti-mouse Alexa Fluor 488, and 8-oxoG cellular density was quantitated. A cellular density threshold was established to define 8-oxoG positive or negative cells. Bars denote mean percentage of 8-oxoG positive cells. Un-paired two-tailed t-test ** p<0.01. N=2 throughout. Error bars=SD of two independent experiments, each performed with three replicates. Representative immunofluorescence images of EP and DS fibroblasts. DAPI (blue), 8-oxoG (green). Scale bar denotes 100μm. **(D)** EP and DS cells were stained with DAPI and mouse anti-BrdU Alexa Fluor 488. Using the secondary only control, a nuclear intensity threshold was established to define BrdU positive or negative nuclei. Bars denote mean percentage of BrdU positive nuclei. Un-paired two-tailed t-test ** p<0.01. N=2 throughout. Error bars, SD of two independent experiments, each performed with three replicates. Representative immunofluorescence images of EP and DS fibroblasts. DAPI (blue), BrdU (green). Digital zoom. Size bar, 50μm.

**Figure S2.**
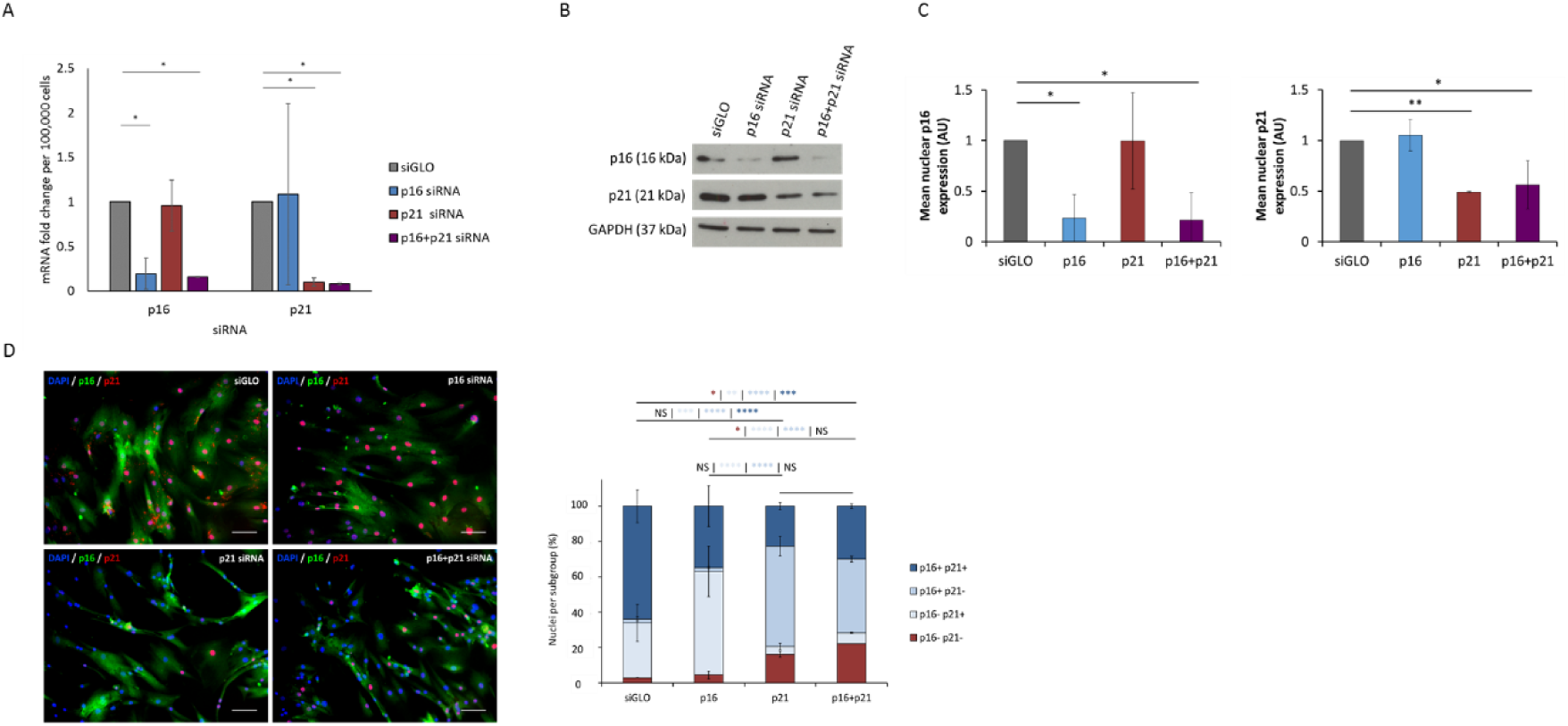
Further confirmation of reversal of deep senescence with an extended panel of markers. **(A-D)** DS HMFs were seeded at 15,000 cells/cm^2^ in 96-well plate format or 384-well plate format and forward transfected with 30nM control siRNA (siGLO), 30nM *p16* siRNA (p16), 30nM *p21* siRNA (p21), or 15nM *p16* siRNA together with 15nM *p21* siRNA (p16+p21) and harvested for RTqPCR at 72 hours post transfection (A) or at five days post transfection for western blot analysis (B-C) or immunofluorescence (D). **(A)** RTqPCR analysis of mRNA levels of *p16* and *p21* in DS HMFs following transfection with control siRNA (siGLO), *p16* siRNA (p16), *p21* siRNA (p21), or *p16* siRNA together with *p21* siRNA (p16+p21). * p<0.05, ** p<0.01. Error bars=SD from two independent experiments, each performed with two replicates. **(B-C)** Western blot depicting p16, p21, and GAPDH levels in DS HMFs following transfection with siGLO, *p16* siRNA, *p21* siRNA, and *p16* in combination with *p21* siRNA (p16+p21 siRNA). Lysates were probed for rabbit anti-p21 (12D1), mouse anti-p16 (JC8), and the rabbit anti-GAPDH antibody. Densitometry analysis of p16 and p21 levels in transfected DS HMFs. * p<0.05, ** p<0.01. Error bars, SD normalised to siGLO siRNA of two independent experiments. **(D)** Nuclear intensity thresholds were established for p16 and p21 to define positive or negative nuclei. Nuclei were classified into four subgroups: p16 and p21 positive (p16+ p21+); p16 positive and p21 negative (p16+ p21-); p16 negative and p21 positive (p16-p21+); and p16 and p21 negative (p16-p21-). Bars denote mean percentage of nuclei per subgroup. Two-way ANOVA and Tukey’s test * p<0.05, ** p<0.01, *** p<0.001, **** p<0.0001. NS=not significant. Significance colours match nuclei subgroup and are ordered left to right in the following order: p16-p21-; p16-p21+; p16+ p21-; p16+ p21+. N=2 throughout. Error bars, SD of two independent experiments, each performed with three replicates. Representative immunofluorescence images of siGLO, *p16, p21*, and *p16*+*p21* siRNA transfected DS HMFs. DAPI (blue), p16 (green), p21 (red). Size bar, 100μm.

**Figure S3.**
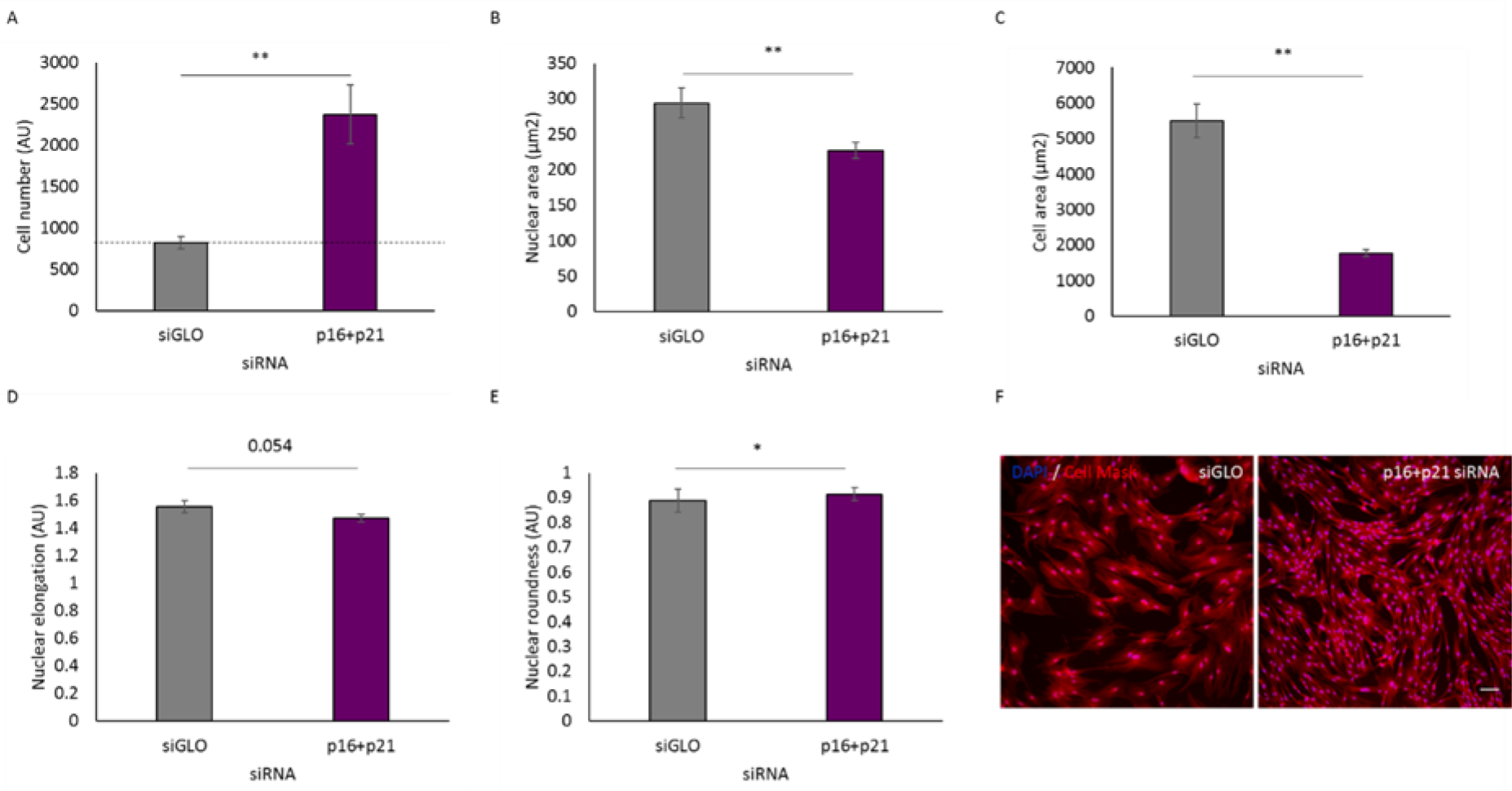
Deep senescence is reversible in primary adult human dermal fibroblasts (HDFs). **(A-F)** DS HDFs were seeded at 10,000 cells/cm^2^ in 384-well plate format and forward transfected with 30nM siGLO (siGLO), 30nM *p16* siRNA (p16), 30nM *p21* siRNA (p21), or 15nM *p16* together with 15nM *p21* siRNA (p16+p21). After five days, cells were fixed, stained with DAPI and Cell Mask, imaged, and quantified. **(A)** Bar chart depicting mean cell number/well for each condition. Dashed line indicates original cell seeded number. ** p<0.01. Error bars, SD from two independent experiments, each performed with three replicates. **(B)** Bar chart depicting mean nuclear area (µm^2^) for each condition. ** p<0.01. Error bars, SD from two independent experiments, each performed with three replicates. **(C)** Bar chart depicting mean cell area (µm^2^) for each condition. ** p<0.01. Error bars, SD from two independent experiments, each performed with three replicates. **(D)** Bar chart depicting mean nuclear elongation (AU) for each condition. Error bars, SD from two independent experiments, each performed with three replicates. **(E)** Bar chart depicting mean nuclear roundness (AU) for each condition. * p<0.05. Error bars, SD from two independent experiments, each performed with three replicates. **(F)** Representative images of DS HDFs stained with DAPI (blue) and Cell Mask (red) following transfection with control siRNA (30nM siGLO) or 15nM *p16* siRNA together with 15nM *p21* siRNA (p16+p21). Size bar, 100µm.

**Figure S4.**
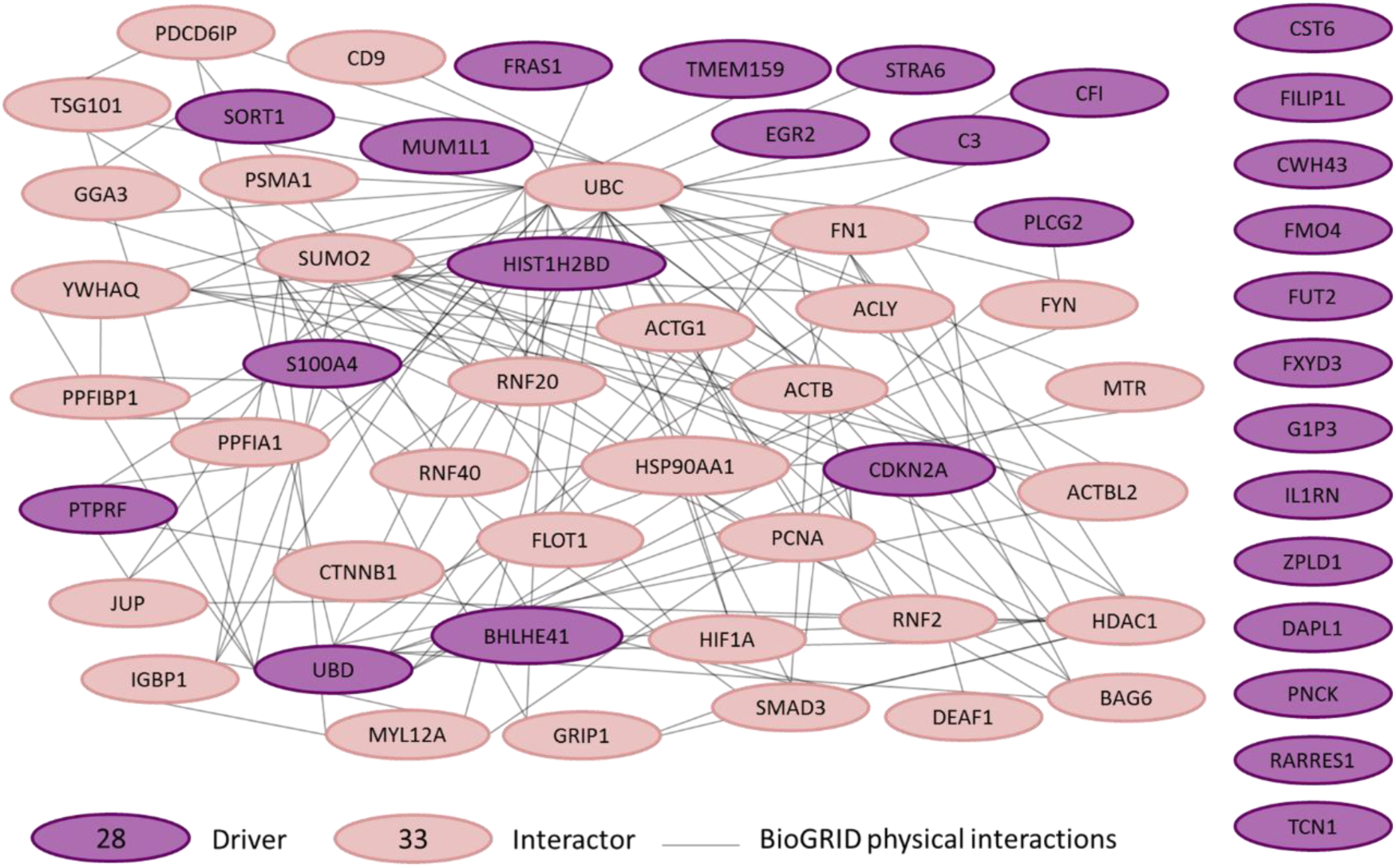
Protein interaction network. Interaction map for twenty-eight potential drivers of senescence (purple) identified in the previous DS HMEC siRNA screen and 33 interactors (pink) identified using bioinformatics to generate a chain with at least two other drivers. Lines represent physical interactions determined from the BioGRID database. Thirteen drivers were not found to be present in this network (listed on the right).

**Figure S5.**
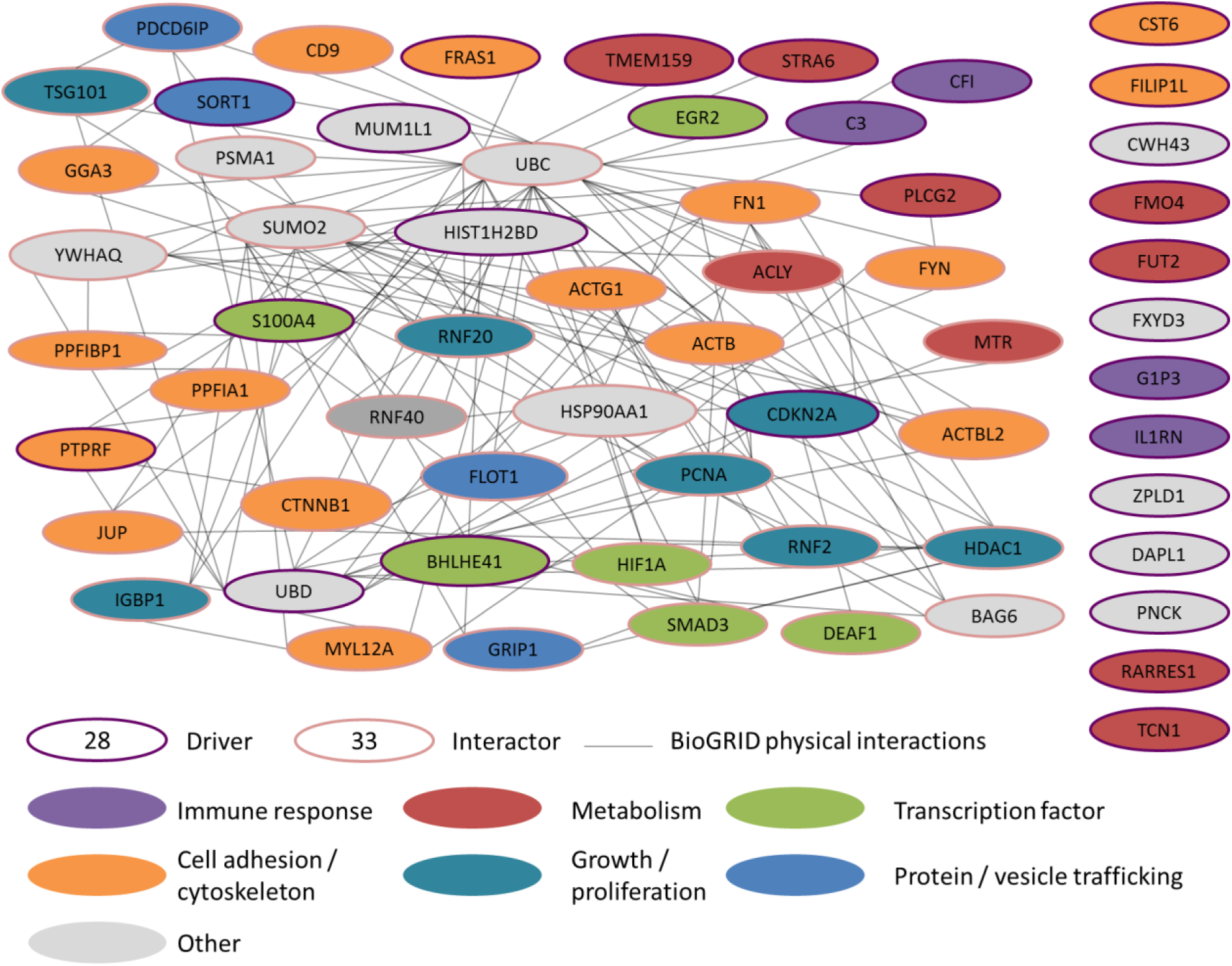
Functional subgrouping of protein interaction network. Using PANTHER, KEGG pathways, and GO bioinformatics tools, potential drivers (outlined in purple) and interactors (outlined in pink) were subgrouped into functional categories: immune response (purple), metabolism (red), transcription factor (green), cell adhesion/cytoskeleton (orange), growth/proliferation (teal), protein/vesicle trafficking (blue), other (grey). Lines represent BioGRID physical interactions.

**Figure S6.**
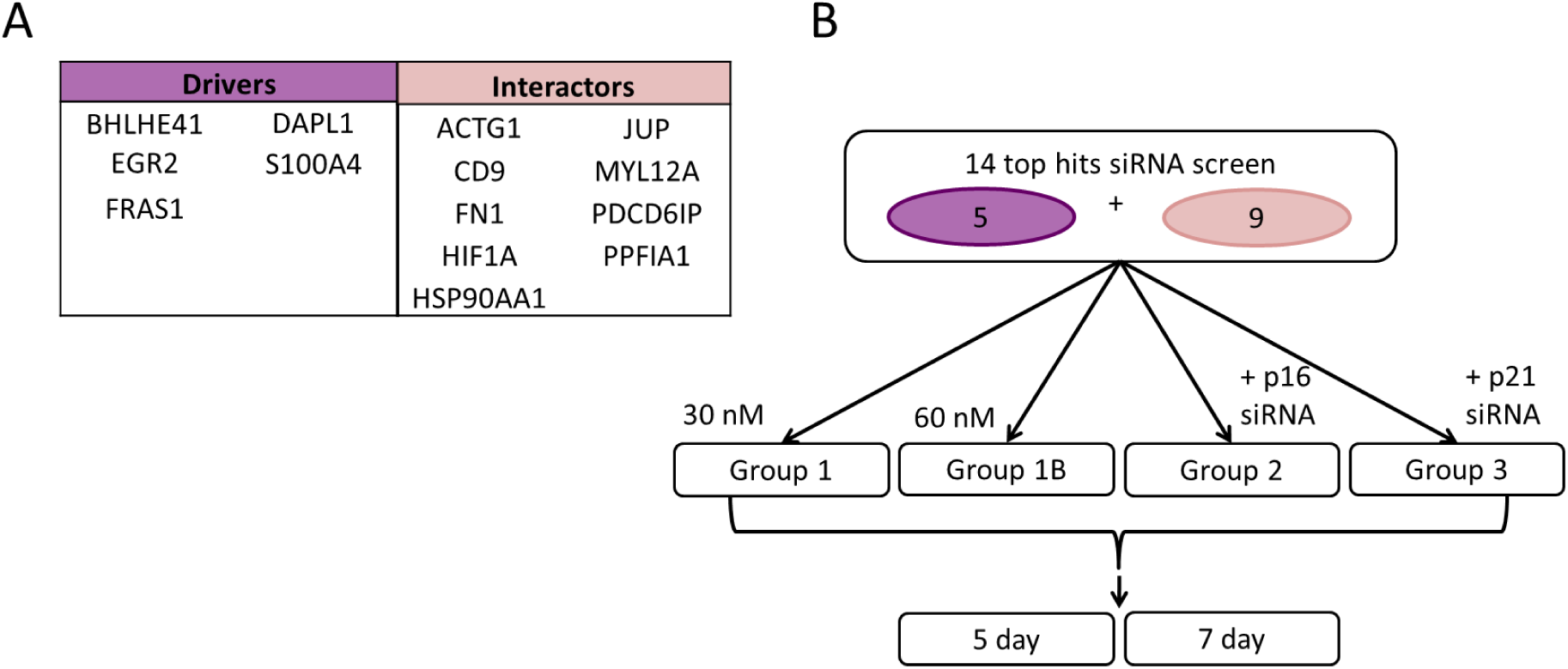
Gene list of the 14 top hits and siRNA screening workflow for investigation of dose effect and time point extension. **(A)** Following the DS HMF screen, the 14 top hits were selected for further investigation: five drivers (purple), and nine interactors (pink). **(B)** Schematic illustrating the experimental design of the smaller siRNA screen investigating dose effect and time point extension in the top 14 hits. Using a previously optimised seeding density and transfection reagent dose, DS HMFs were forward transfected with the 14 siRNAs in four conditions: 30nM siRNA individually (Group 1); 60nM siRNA individually (Group 1B); 15nM siRNA in combination with 15nM siRNA (Group 2); and 15nM siRNA in combination with p21 siRNA (Group 3). After five or seven days, cells were fixed and stained with DAPI and Cell Mask, and quantified.

**Figure S7.**
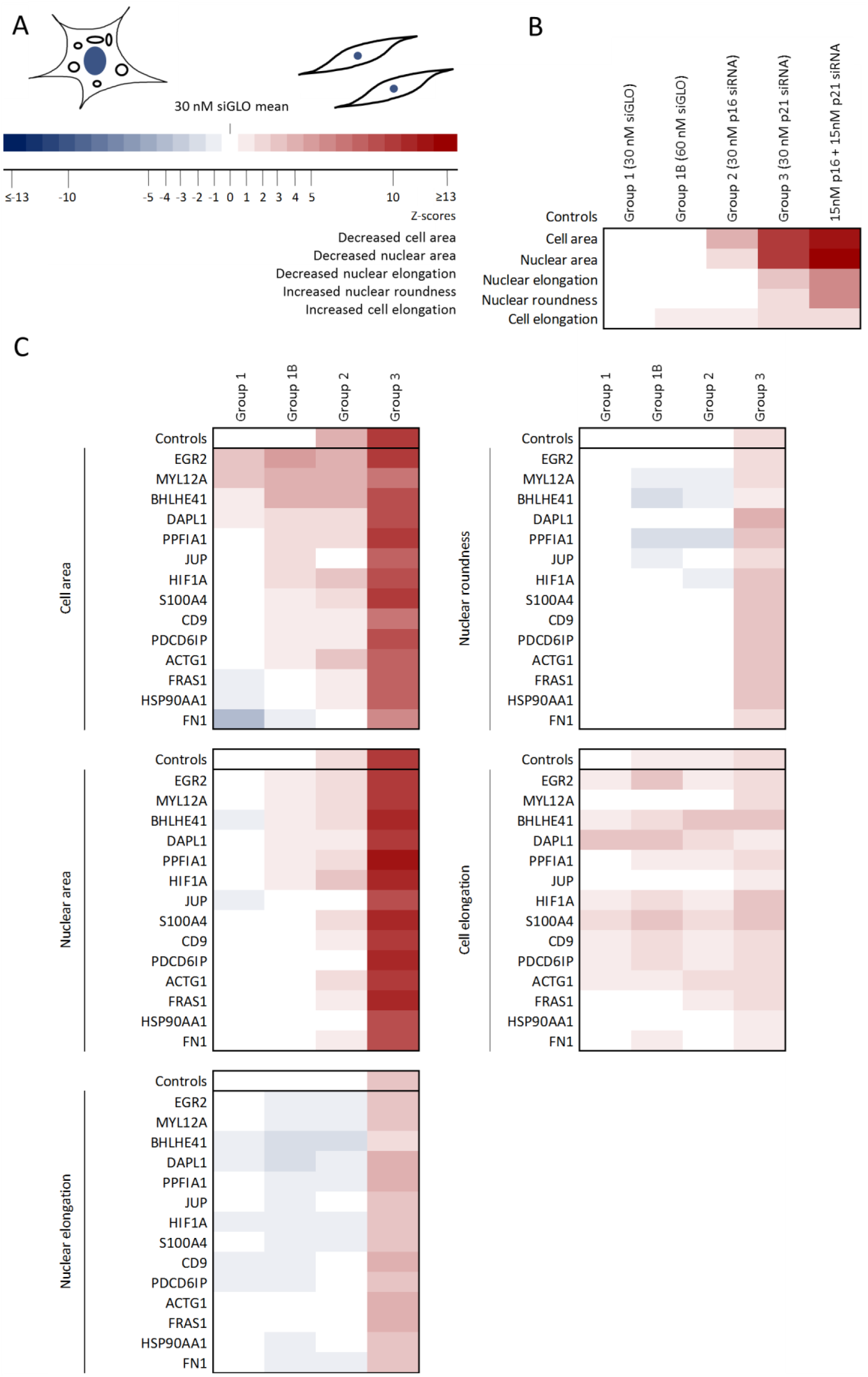
Multi-parameter analysis of the top 14 siRNAs identified to reverse senescence in the DS HMFs at day five. Three independent siRNA screens, each in triplicate, were performed for the five day time point in DS HMFs. Cells were fixed, stained with DAPI and Cell Mask, imaged and nuclear and cellular morphologies quantified. Z scores were then generated. **(A)** Key. The colour saturation reflects the number of Z scores from the siGLO control mean. Scores highlighted in red denote a shift towards the reversed phenotype and blue denotes a shift away from the reversed phenotype. **(B)** Heatmap depicting significant changes in each of the panel of five morphological senescence-associated markers from the 30nM siGLO control mean (Group 1 control) for 60nM siGLO (Group 1B control), 30nM p16 siRNA (p16) (Group 2 control), 30nM p21 siRNA (p21) (Group 3 control), and 15nM p16 together with 15nM p21 siRNA (p16+p21) transfected DS HMFs. **(C)** Heatmap depicting significant changes in each of the panel of five morphological senescence-associated markers for the hit siRNAs selected from the previous screen in four different conditions: 30nM siRNA individually (Group 1); 60nM siRNA individually (Group 1B); 15nM siRNA in combination with 15nM siRNA (Group 2); and 15nM siRNA in combination with p21 siRNA (Group 3), compared to 30nM siGLO control mean.

**Figure S8.**
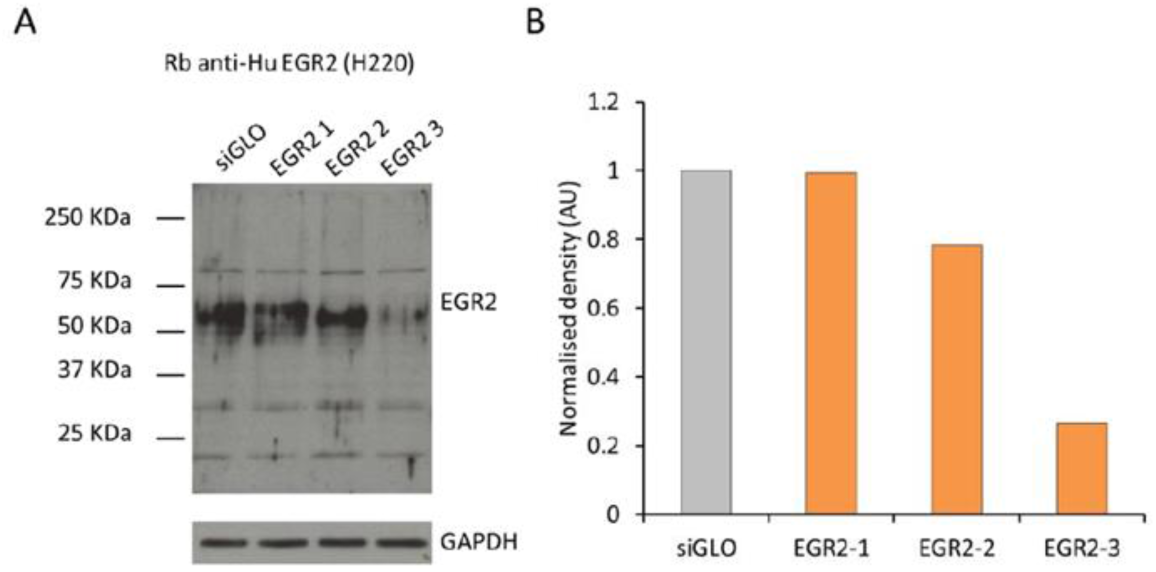
Uncropped western blot of rabbit anti-human EGR2 (H220) antibody in DS HMFs transfected with individual EGR2 siRNAs. **(A)** Representative western blot of DS HMFs transfected with 30nM siGLO (siGLO) or 30nM individual EGR2 siRNA (‘*EGR2 1*’, ‘*EGR2 2*’, *‘EGR2 3’* cell lysates probed for rabbit anti-human EGR2 (H220). **(B)** Densitometry analysis of EGR2 bands in rabbit anti-human EGR2 (H220) probed transfected DS HMFs. Analysis was performed using ImageJ software. Bars denote density levels normalised to GAPDH relative to the siGLO control.

**Figure S9.**
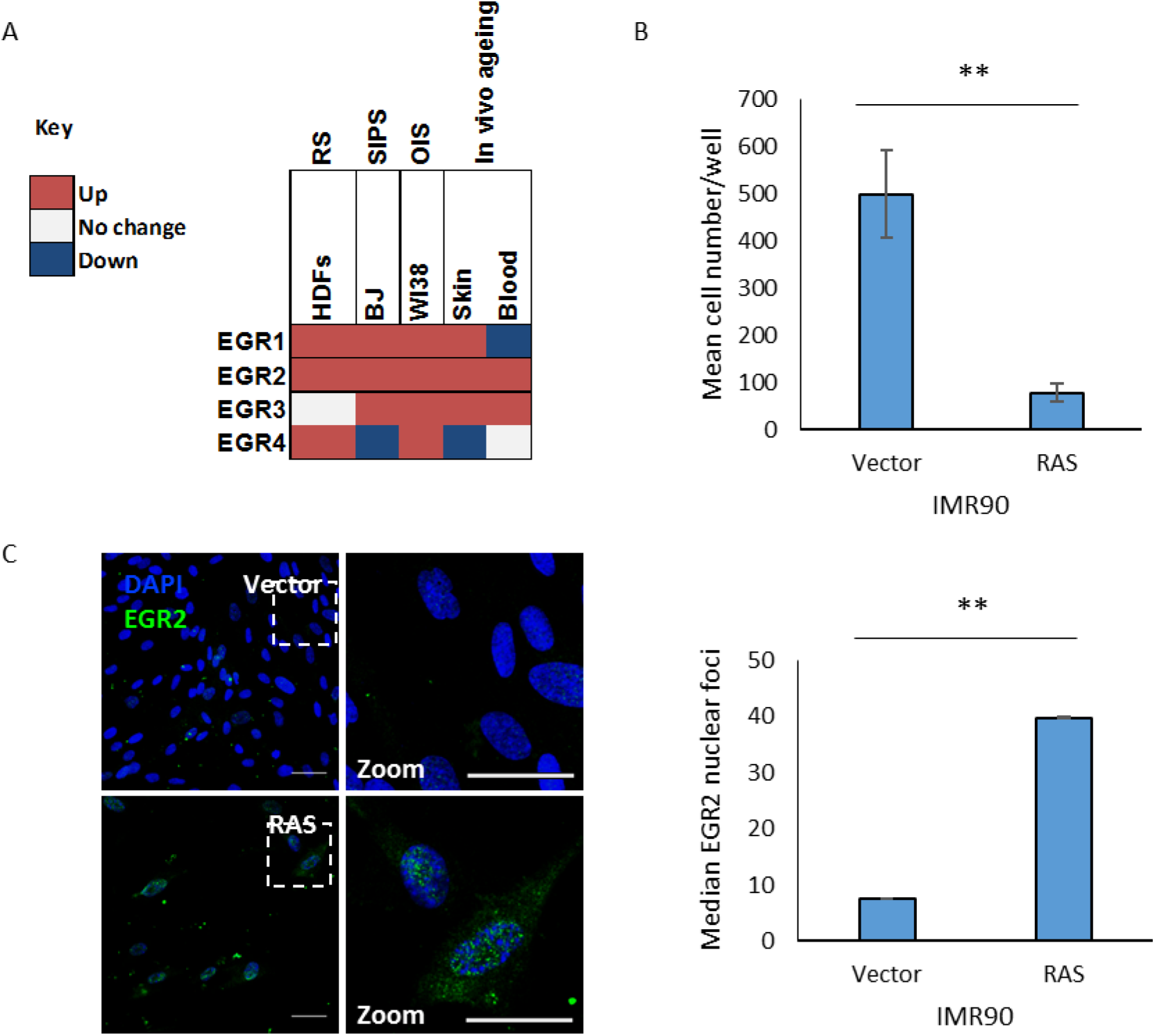
EGR2 expression increases in multiple models of senescence *in vitro*, human ageing *in vivo*, and EGR2 protein levels increase in RAS oncogene-induced senescence (OIS) fibroblasts. **(A)** EGR2 gene expression during *in vitro* senescence for replicatively senescent (RS) primary adult dermal fibroblasts (HDFs), bleomycin-induced stress-induced premature senescence (SIPS), RAS OIS in WI38 foetal lung fibroblasts, and *in vivo* ageing of human skin and blood. Red indicates an increase and blue indicates a decrease in EGR2 expression. **(B)** Barchart depicting mean cell number per well for vector and RAS OIS fibroblasts (RAS). ** p<0.01. Error bars, SD from three independent experiments, each performed with two replicates. **(C)** Representative immunofluorescence images of vector and RAS OIS fibroblasts stained with DAPI (blue) and EGR2 (green) at 5 days post-seeding. Barchart depicting median EGR2 nuclear foci in vector and RAS OIS fibroblasts (RAS). ** p<0.01. Error bars, SD from three independent experiments, each performed with two replicates.

**Figure S10.**
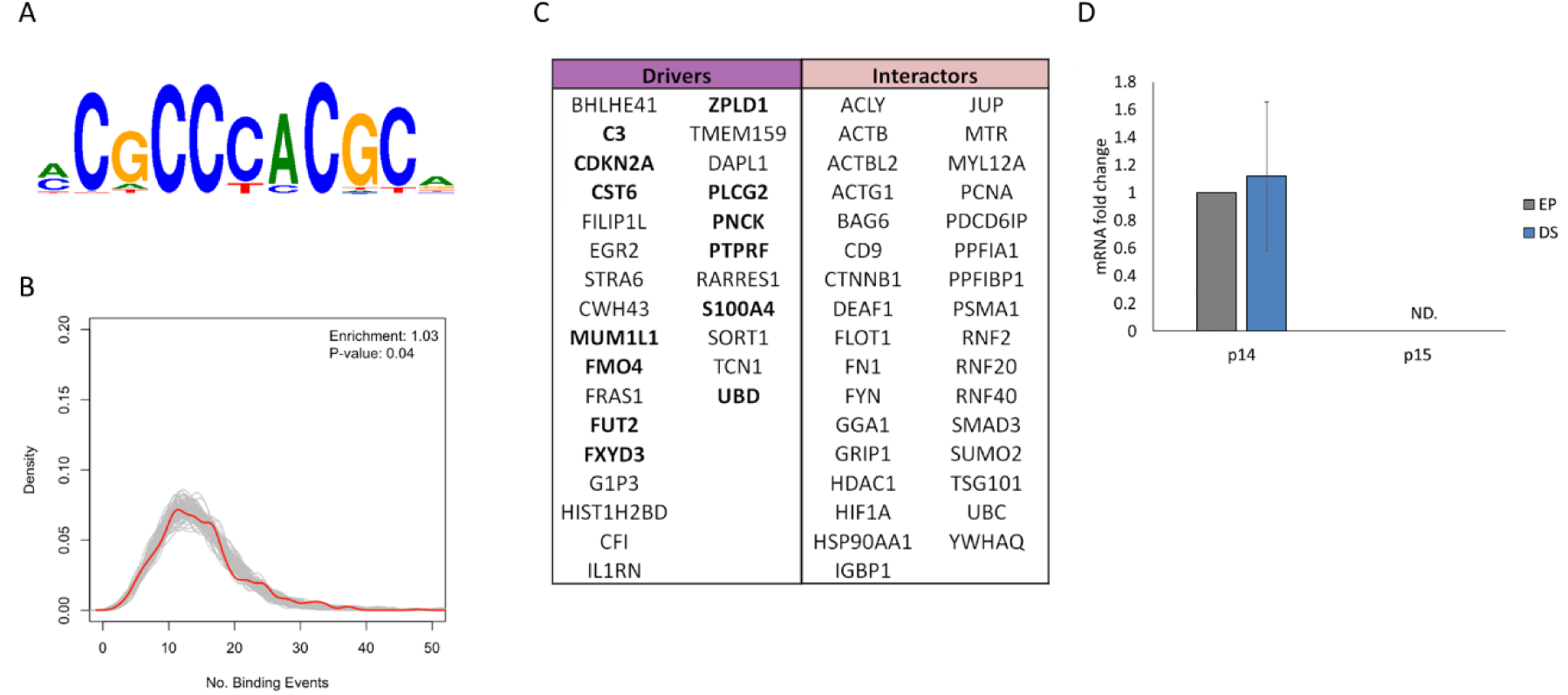
Predicted EGR2 binding sites within siRNA screening hits and INK4/ARF locus. **(A)** EGR2 DNA binding motif (Jolma *et al*., 2013; Mathelier *et al*., 2016). **(B)** The EGR2 DNA binding motif was investigated in the promoters of genes up-regulated in HMEC senescence (red) relative to random sampling (grey). **(C)** The gene promoters predicted to contain EGR2 DNA binding motifs in the DS HMF siRNA screen are presented here in bold. **(D)** RTqPCR analysis of mRNA levels of *p14 (ARF)* and *p15* in EP and DS HMFs. Error bars, SD from two independent experiments, each performed with two replicates.

**Figure S11.**
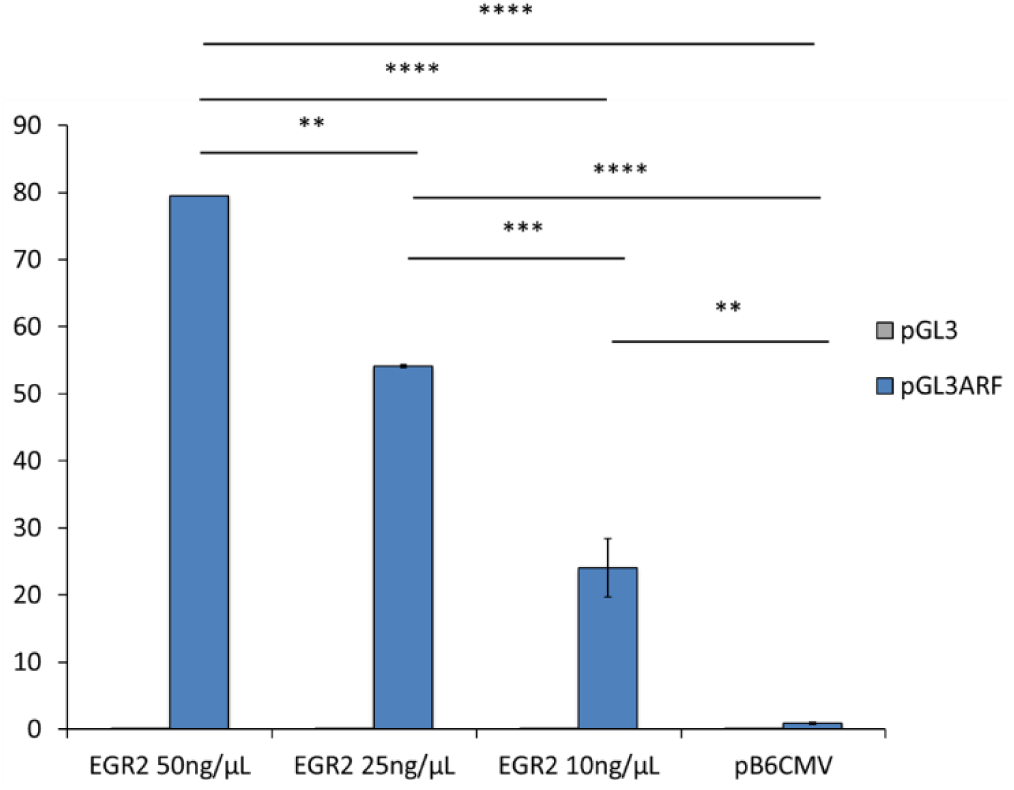
EGR2 expression vector titration results in ARF activation dose response. Mean luciferase values for activation of pGL3 luciferase reporter constructs with and without the *ARF* promoter sequence (pGL3 or pGL3 ARF, respectively) following co-transfection of U2OS cells with titratable amounts of expression vector encoding EGR2 (50ng/µL, 25ng/µL, 10ng/µL, respectively) compared to pB6CMV vector backbone. ** p<0.01, *** p<0.001, **** p<0.0001. Error bars, SD from two experiments.

**Figure S12.**
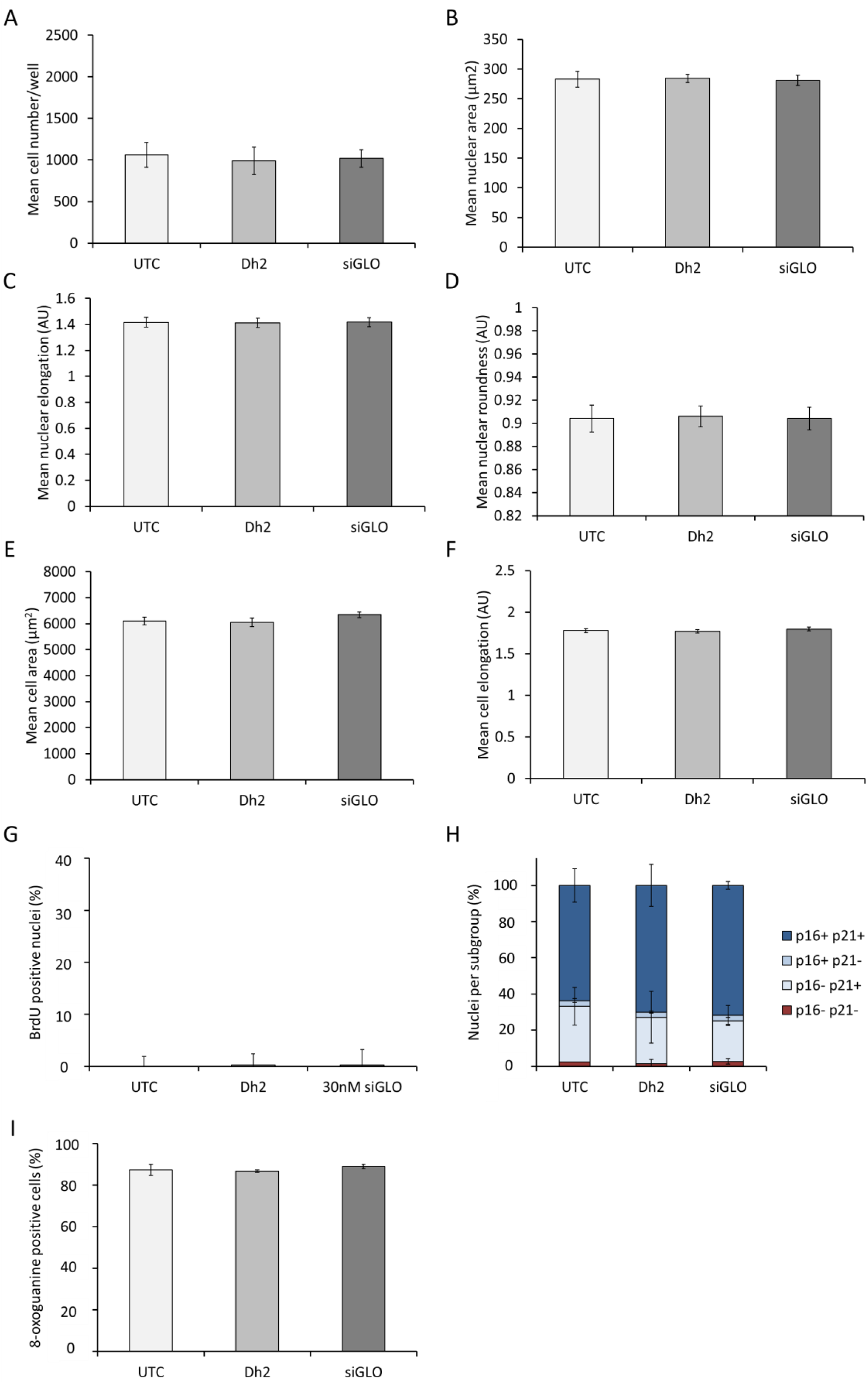
Analysis of untransfected, transfection reagent only, and siGLO transfected controls. **(A-I)** DS HMFs were seeded at 15,000 cells/cm^2^ in 384-well plate format and were either untransfected (UTC) or forward transfected with either transfection reagent alone (Dh2), or with 30nM siGLO (siGLO). Cells were then fixed, stained with DAPI, mouse anti-BrdU Alexa Fluor 488, or mouse anti-p16 (JC8) and rabbit anti-p21 (12D1), or mouse anti-8-oxoguanine, and donkey anti-mouse Alexa Fluor 488 and goat anti-rabbit Alexa Fluor 546, and Cell Mask. Nuclear and cellular morphologies, BrdU nuclear intensity, p16 and p21 nuclear intensities, and 8-oxoguanine cellular density was quantitated. Using the secondary only control, a nuclear intensity threshold was established to define BrdU positive or negative nuclei. Nuclear intensity thresholds were established for p16 and p21 to define positive or negative nuclei. Nuclei were classified into four subgroups: p16 and p21 positive (p16+ p21+); p16 positive and p21 negative (p16+ p21-); p16 negative and p21 positive (p16-p21+); and p16 and p21 negative (p16-p21-). A cellular density threshold was established to define 8-oxoguanine positive or negative cells. **(A)** Bar chart depicting mean cell number/well. N=4. Error bars=SD of four independent experiments, each performed with three replicates. **(B)** Bar chart depicting mean nuclear area (μm^2^). N=4. Error bars=SD of four independent experiments, each performed with three replicates. **(C)** Bar chart depicting mean nuclear elongation (μm^2^). N=4. Error bars=SD of four independent experiments, each performed with three replicates. **(D)** Bar chart depicting mean nuclear roundness (AU). N=4. Error bars=SD of four independent experiments, each performed with three replicates. **(E)** Bar chart depicting mean cell area (μm^2^). N=4. Error bars=SD of four independent experiments, each performed with three replicates. **(F)** Bar chart depicting mean cell elongation (AU). N=4. Error bars=SD of four independent experiments, each performed with three replicates. **(G)** Bars denote mean percentage of BrdU positive nuclei. N=1. Error bars=SD of a single experiment containing three replicates. **(H)** Bars denote mean percentage of nuclei per subgroup. N=2. Error bars=SD of two independent experiments, each performed with three replicates. **(I)** Bars denote mean percentage of 8-oxoguanine positive nuclei. N=2. Error bars=SD of two independent experiments, each performed with three replicates.

## Notes

### Competing Interest Statement

The authors have declared no competing interest.

## References

Alcorta, D. A. et al. (1996) ‘Involvement of the cyclin-dependent kinase inhibitor p16 (INK4a) in replicative senescence of normal human fibroblasts.’, Proceedings of the National Academy of Sciences of the United States of America, 93(24), pp. 13742–7. doi: 10.1073/pnas.93.24.13742.

del Arroyo, A. G. et al. (2007) ‘E2F-Dependent Induction of p14ARF During Cell Cycle Re-Entry in Human T Cells’, Cell Cycle, 6(21), pp. 2697–2705. doi: 10.4161/cc.6.21.4857.

Atanasoski, S. et al. (2006) ‘Cell cycle inhibitors p21 and p16 are required for the regulation of Schwann cell proliferation’, Glia, 53(2), pp. 147–157. doi: 10.1002/glia.20263.

Baker, D. J. et al. (2011) ‘Clearance of p16Ink4a-positive senescent cells delays ageing-associated disorders.’, Nature. Nature Publishing Group, 479(7372), pp. 232–6. doi: 10.1038/nature10600.

Baker, D. J. et al. (2016) ‘Naturally occurring p16 Ink4a -positive cells shorten healthy lifespan’, Nature. Nature Publishing Group, 530(7589), pp. 1–5. doi: 10.1038/nature16932.

Beauséjour, C. M. et al. (2003) ‘Reversal of human cellular senescence: Roles of the p53 and p16 pathways’, EMBO Journal, 22(16), pp. 4212–4222. doi: 10.1093/emboj/cdg417.

Beckmann, A. M. and Wilce, P. A. (1997) ‘Egr transcription factors in the nervous system’, Neurochemistry International, 31(4), pp. 477–510. doi: 10.1016/S0197-0186(97)00001-6.

Bishop, C. L. et al. (2010) ‘Primary cilium-dependent and -independent hedgehog signaling inhibits p16INK4A’, Molecular Cell. Elsevier Inc., 40(4), pp. 533–547. doi: 10.1016/j.molcel.2010.10.027.

Borgdorff, V. et al. (2010) ‘Multiple microRNAs rescue from Ras-induced senescence by inhibiting p21(Waf1/Cip1)’, Oncogene. Nature Publishing Group, 29(15), pp. 2262–2271. doi: 10.1038/onc.2009.497.

Brummelkamp, T. R. et al. (2002) ‘TBX-3, the Gene Mutated in Ulnar-Mammary Syndrome, Is a Negative Regulator of p19ARF and Inhibits Senescence*’, The Journal of Biological Chemistry, 277(8), pp. 6567–6572. doi: 10.1074/jbc.M110492200.

Carvalho, C. et al. (2019) ‘Glucocorticoids delay RAF-induced senescence promoted by EGR1’, Journal of Cell Science, 132(16), p. jcs230748. doi: 10.1242/jcs.179960.

Coppé, J.-P. et al. (2008) ‘Senescence-associated secretory phenotypes reveal cell-nonautonomous functions of oncogenic RAS and the p53 tumor suppressor.’, PLoS biology, 6(12), pp. 2853–2868. doi: 10.1371/journal.pbio.0060301.

Decker, L. (2006) ‘Peripheral Myelin Maintenance Is a Dynamic Process Requiring Constant Krox20 Expression’, Journal of Neuroscience, 26(38), pp. 9771–9779. doi: 10.1523/JNEUROSCI.0716-06.2006.

Delmas, V. et al. (2007) ‘β-Catenin induces immortalization of melanocytes by suppressing’, Genes & Development, 21, pp. 2923–2935. doi: 10.1101/gad.450107.al.

van Deursen, J. M. (2014) ‘The role of senescent cells in ageing.’, Nature. Nature Publishing Group, 509(7501), pp. 439–46. doi: 10.1038/nature13193.

Dimri, G. P. et al. (2000) ‘Regulation of a Senescence Checkpoint Response by the E2F1 Transcription Factor and p14 ARF Tumor Suppressor’, Molecular and Cellular Biology, 20(1), pp. 273–285.

Douville, J. M. et al. (2011) ‘Mechanisms of MEOX1 and MEOX2 Regulation of the Cyclin Dependent Kinase Inhibitors p21 CIP1 / WAF1 and p16 INK4a in Vascular Endothelial Cells’, PLoS ONE, 6(12), pp. 1–16. doi: 10.1371/journal.pone.0029099.

Dyson, N. (1998) ‘The regulation of E2F by pRB-family?proteins’, Genes & Development, 12(617), pp. 2245–2262. doi: 10.1101/gad.12.15.2245.

Freund, A. et al. (2012) ‘Lamin B1 loss is a senescence-associated biomarker.’, Molecular biology of the cell, 23(11), pp. 2066–75. doi: 10.1091/mbc.E11-10-0884.

Garbe, J. C. et al. (2009) ‘Molecular distinctions between stasis and telomere attrition senescence barriers shown by long-term culture of normal human mammary epithelial cells.’, Cancer research, 69, pp. 7557–7568. doi: 10.1158/0008-5472.CAN-09-0270.

Garbe, J. C. et al. (2014) ‘Immortalization of normal human mammary epithelial cells in two steps by direct targeting of senescence barriers does not require gross genomic alterations’, Cell Cycle, 13(21), pp. 3423–3435. doi: 10.4161/15384101.2014.954456.

Gil, J. et al. (2004) ‘Polycomb CBX7 has a unifying role in cellular lifespan.’, Nature cell biology, 6, pp. 67–72. doi: 10.1038/ncb1077.

Gil, J. and Peters, G. (2006) ‘Regulation of the INK4b-ARF-INK4a tumour suppressor locus: all for one or one for all.’, Nature Reviews. Molecular cell biology, 7, pp. 667–677. doi: 10.1038/nrm1987.

Gire, V. and Wynford-Thomas, D. (1998) ‘Reinitiation of DNA synthesis and cell division in senescent human fibroblasts by microinjection of anti-p53 antibodies.’, Molecular and cellular biology, 18(3), pp. 1611–21.

Hari, P. et al. (2019) ‘The innate immune sensor Toll-like receptor 2 controls the senescence-associated secretory phenotype’, Science Advances, 5(6), pp. 1–15. doi: 10.1126/sciadv.aaw0254.

Hayflick, L. and Moorhead, P. S. (1961) ‘The serial cultivation of human diploid cell strains.’, Journal of Chemical Information and Modeling, 53(9), pp. 1689–1699. doi: 10.1017/CBO9781107415324.004.

Irelan, J. T. et al. (2009) ‘A functional screen for regulators of CKDN2A reveals MEOX2 as a transcriptional activator of INK4a’, PLoS ONE, 4(4), pp. 2–9. doi: 10.1371/journal.pone.0005067.

Jacobs, J. J. L. et al. (2000) ‘Senescence bypass screen identifies TBX2, which represses Cdkn2a (p19 ARF) and is amplified in a subset of human breast cancers’, Nature Genetics, 26(November), pp. 291–299.

Jolma, A. et al. (2013) ‘DNA-binding specificities of human transcription factors’, Cell. Elsevier Inc., 152(1–2), pp. 327–339. doi: 10.1016/j.cell.2012.12.009.

Katayama, K. et al. (2008) ‘FOXO transcription factor-dependent p15 INK4b and p19 INK4d expression’, Oncogene, 27, pp. 1677–1686. doi: 10.1038/sj.onc.1210813.

Kim, You-mi et al. (2013) ‘Implications of time-series gene expression profiles of replicative senescence’, Aging Cell, 12, pp. 622–634. doi: 10.1111/acel.12087.

Krishnamurthy, J. et al. (2006) ‘p16INK4a induces an age-dependent decline in islet regenerative potential’, Nature, 443(7110), pp. 453–457. doi: nature05092 [pii] 10.1038/nature05092.

Krones-Herzig, A., Adamson, E. and Mercola, D. (2003) ‘Early growth response 1 protein, an upstream gatekeeper of the p53 tumor suppressor, controls replicative senescence.’, Proceedings of the National Academy of Sciences of the United States of America, 100(6), pp. 3233–3238. doi: 10.1073/pnas.2628034100.

Lowe, R. et al. (2015) ‘The senescent methylome and its relationship with cancer, ageing and germline genetic variation in humans.’, Genome biology. Genome Biology, 16, p. 194. doi: 10.1186/s13059-015-0748-4.

Martin, N. et al. (2013) ‘Co-regulation of senescence-associated genes by oncogenic homeobox proteins and polycomb repressive complexes’, Cell Cycle, 12(January 2015), pp. 2194–2199. doi: 10.4161/cc.25331.

Martin, N., Beach, D. and Gil, J. (2014) ‘Ageing as developmental decay: insights from p16INK4a’, Trends in Molecular Medicine, 20(12), pp. 667–674. doi: 10.1016/j.molmed.2014.09.008.

Martínez-Zamudio, R. I. et al. (2020) ‘AP-1 imprints a reversible transcriptional programme of senescent cells’, Nature Cell Biology. doi: 10.1038/s41556-020-0529-5.

Mathelier, A. et al. (2016) ‘JASPAR 2016: A major expansion and update of the open-access database of transcription factor binding profiles’, Nucleic Acids Research, 44(D1), pp. D110–D115. doi: 10.1093/nar/gkv1176.

Naylor, R. M., Baker, D. J. and van Deursen, J. M. (2012) ‘Senescent Cells: A Novel Therapeutic Target for Aging and Age-Related Diseases’, Clinical Pharmacology & Therapeutics, 93(1), pp. 105–116. doi: 10.1038/clpt.2012.193.

Ohtani, N. et al. (2001) ‘Opposing effects of Ets and Id proteins on p16INK4a expression during cellular senescence.’, Nature, 409(6823), pp. 1067–1070. doi: 10.1038/35059131.

Parkinson, D. B. et al. (2004) ‘Krox-20 inhibits Jun-NH2-terminal kinase/c-Jun to control Schwann cell proliferation and death’, Journal of Cell Biology, 164, pp. 385–394. doi: 10.1083/jcb.200307132.

Passos, J. F. et al. (2010) ‘Feedback between p21 and reactive oxygen production is necessary for cell senescence.’, Molecular systems biology, 6(347), p. 347. doi: 10.1038/msb.2010.5.

Peters, M. J. et al. (2015) ‘The transcriptional landscape of age in human peripheral blood.’, Nature communications, 6, p. 8570. doi: 10.1038/ncomms9570.

Rodier, F. et al. (2009) ‘Persistent DNA damage signaling triggers senescence-associated inflammatory cytokine secretion’, Nature Cell Biology, 11(8), pp. 973–979. doi: 10.1038/ncb1909.Persistent.

Romanov, S. R. et al. (2001) ‘Normal human mammary epithelial cells spontaneously escape senescence and acquire genomic changes.’, Nature, 409(6820), pp. 633–637. doi: 10.1038/35054579.

šafka Brožková, D. et al. (2012) ‘Charcot-Marie-Tooth neuropathy due to a novel EGR2 gene mutation with mild phenotype - Usefulness of human mapping chip linkage analysis in a Czech family’, Neuromuscular Disorders, 22, pp. 742–746. doi: 10.1016/j.nmd.2012.04.002.

Serrano, M. et al. (1997) ‘Oncogenic ras provokes premature cell senescence associated with accumulation of p53 and p16(INK4a)’, Cell, 88, pp. 593–602. doi: 10.1016/S0092-8674(00)81902-9.

Sharpless, N. E. and Sherr, C. J. (2015) ‘Forging a signature of in vivo senescence’, Nature Reviews Cancer. Nature Publishing Group, 15(7), pp. 397–408. doi: 10.1038/nrc3960.

Srinivasan, R. et al. (2012) ‘Genome-wide analysis of EGR2/SOX10 binding in myelinating peripheral nerve’, Nucleic Acids Research, 40(14), pp. 6449–6460. doi: 10.1093/nar/gks313.

Topilko, P. et al. (1994) ‘Krox-20 controls myelination in the peripheral nervous system.’, Nature, pp. 796–799. doi: 10.1038/371796a0.

Unoki, M. and Nakamura, Y. (2003) ‘Methylation at CpG islands in intron 1 of EGR2 confers enhancer-like activity’, FEBS Letters, 554(1–2), pp. 67–72. doi: 10.1016/S0014-5793(03)01092-5.

Wassermann, S. et al. (2009) ‘p16INK4a Is a β-Catenin Target Gene and Indicates Low Survival in Human Colorectal Tumors’, Gastroenterology. AGA Institute American Gastroenterological Association, 136(1), pp. 196-205.e2. doi: 10.1053/j.gastro.2008.09.019.

Yalcin, S. et al. (2008) ‘Foxo3 Is Essential for the Regulation of Ataxia Telangiectasia Mutated and Oxidative Stress-mediated Homeostasis of Hematopoietic Stem Cells * □’, The Journal of Biological Chemistry, 283(37), pp. 25692–25705. doi: 10.1074/jbc.M800517200.

Zhang, Y., Xiong, Y. and Yarbrough, W. G. (1998) ‘ARF Promotes MDM2 Degradation and Stabilizes p53: ARF-INK4a Locus Deletion Impairs Both the Rb and p53 Tumor Suppression Pathways’, Cell, 92, pp. 725–734.

Zheng, Y. et al. (2013) ‘Egr2-dependent gene expression profiling and ChIP-Seq reveal novel biologic targets in T cell anergy.’, Molecular immunology. Elsevier Ltd, 55(3–4), pp. 283–91. doi: 10.1016/j.molimm.2013.03.006.

Zorick, T. S. et al. (1996) ‘The transcription factors SCIP and Krox-20 mark distinct stages and cell fates in Schwann cell differentiation.’, Molecular and cellular neurosciences, 8(2–3), pp. 129–145. doi: 10.1006/mcne.1996.0052.

